# A comprehensive spatiotemporal map of dystrophin isoform expression in the developing and adult human brain

**DOI:** 10.1101/2024.12.20.629620

**Authors:** Francesco Catapano, Reem Alkharji, Darren Chambers, Simran Singh, Arta Aghaeipour, Jyoti Malhotra, Patrizia Ferretti, Rahul Phadke, Francesco Muntoni

## Abstract

Mutations in the dystrophin gene (*DMD)* cause the severe muscle-wasting disease Duchenne Muscular Dystrophy (DMD). Additionally, there is a high incidence of intellectual disability and neurobehavioural comorbidities in individuals with DMD. Similar behavioural abnormalities are found in *mdx* dystrophic mouse models. Unlike muscle, several dystrophin isoforms are expressed in the human brain, but a detailed map of regional and cellular localisation of dystrophin isoforms is missing. This is crucial in understanding the neuropathology of DMD individuals, and for evaluating the translatability of pre-clinical findings in DMD mouse models receiving genetic therapy interventions. Here, we provide a comprehensive dystrophin expression profile in human brains from early development to adulthood. We reveal expression of *dp427p2*, *dp427c*, *dp427m* and *dp40* isoforms in embryonic brains, not previously reported. *Dp427p2* and *dp140* were greatly downregulated in adult brains, although the latter continued to be expressed across several regions. Importantly, we demonstrate for the first-time expression of *DMD* transcripts in human motor neurons and co-expression of different dystrophin isoforms within single neurons in both developing and adult brains. Finally, we show localisation of *DMD* transcripts with *GAD1*+ GABAergic-associated transcripts in neurons including cerebellar Purkinje cells and interneurons, as well as in the majority of neocortical and hippocampal *SLC17A7*+ glutamatergic neurones, suggesting a role for dystrophin in signalling at the neuronal inhibitory and excitatory synapses.

**Graphical Abstract:** 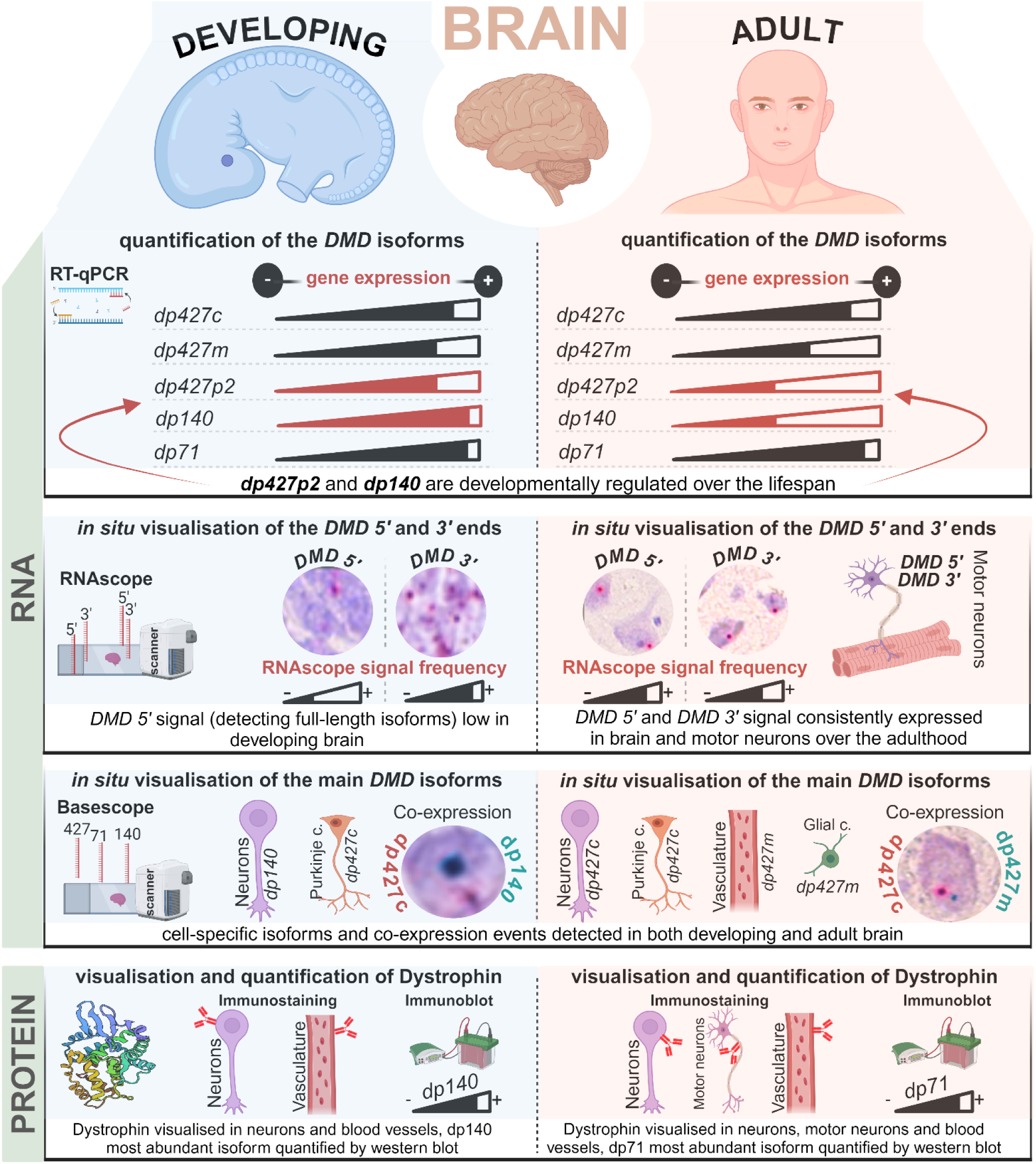

## Introduction

Dystrophin *(DMD)* is the largest gene in the human genome. It consists of 79 exons located on the X chromosome and produces multiple transcripts, whose expression is regulated in a cell-tissue-specific fashion by seven promoters [1, 2]. In addition, a set of splice variants is synthesised under the regulation of multiple splicing elements and polyA sites at the *DMD* locus [2] (**Fig. 1**).

**Fig. 1.**
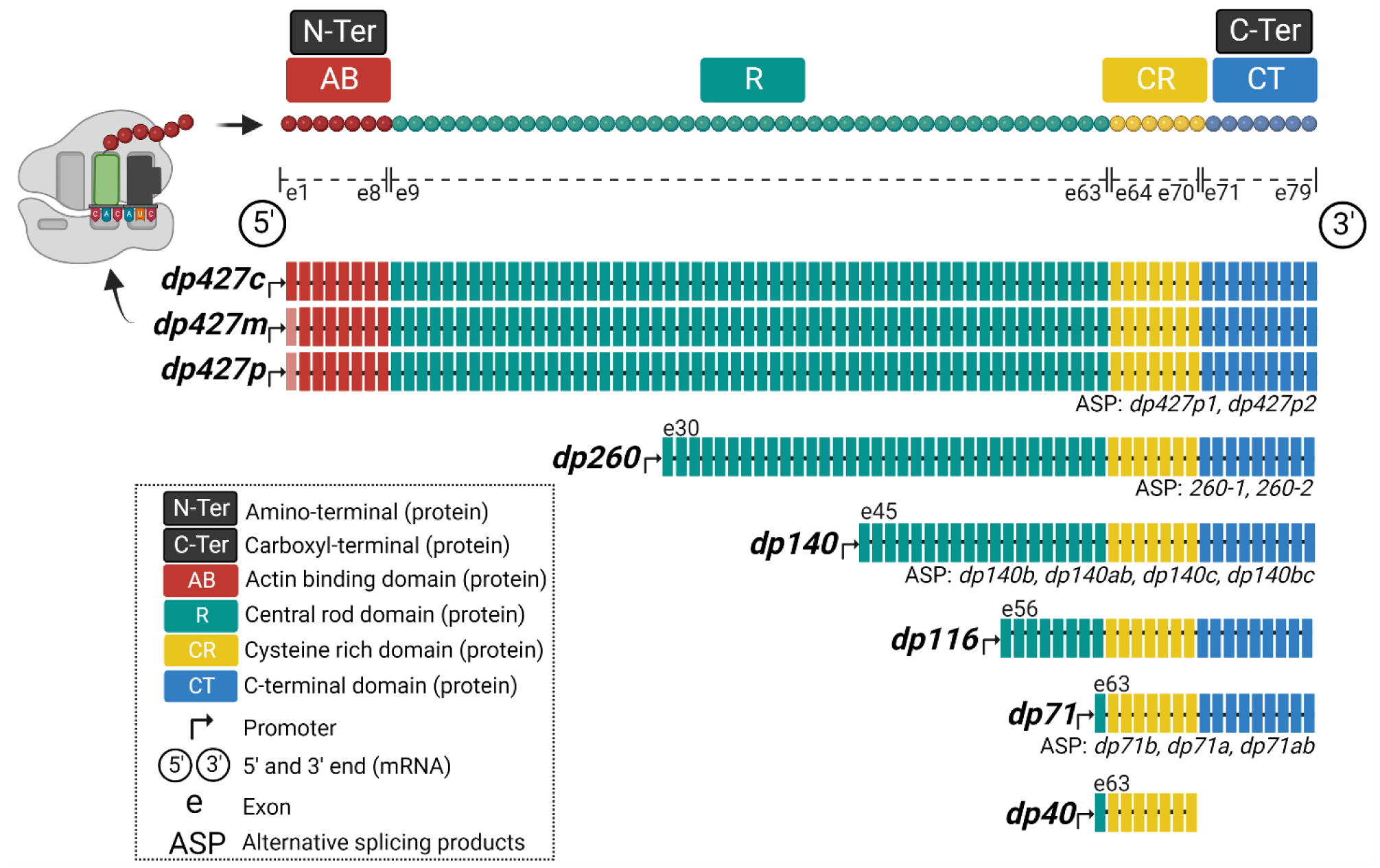
Correlation of human *DMD* transcripts and dystrophin exon-domain. Isoforms derived from the *DMD* gene can be classified into full-, intermediate- and short-length according to their mass in kDa and are all expressed in the human nervous system. *DMD* exons are shown with colours matching the encoded dystrophin functional domains. ASP (alternative splicing products) are generated by alternative splicing and distinct polyA-addition signals.

Therefore, several dystrophin protein isoforms can be produced, and they are classified according to their molecular masses as full-length (*dp427m*, muscle; *dp427c*, cerebral; *dp427p*, Purkinje cells), intermediate-length (*dp260*; *dp140*; *dp116,* mainly expressed in retina, brain and Schwann cells, respectively) and short-length (*dp71* and *dp40,* both more ubiquitously expressed) [3].

Loss of the muscle-specific *dp427m* isoform is the cause of the rare muscle-wasting disorder, Duchenne Muscular Dystrophy (DMD, OMIM #310200), that affects 1 in about 3500 live male births, and results in progressive muscle degeneration leading to premature death because of heart and pulmonary failure [4, 5]. Hence the role of dystrophin, and particularly dp427m, has been extensively investigated and is best understood in muscle [6–8].

However, it is becoming more and more recognised that DMD patients (50-70%) display central nervous system (CNS) co-morbidities, and dystrophin isoforms have been shown to be expressed in the brain [9–16].

It has been suggested that intellectual disability and behavioural comorbidities in DMD patients, including autism spectrum disorder, attention-deficit/hyperactivity disorder and epilepsy, depend on the *DMD* mutation site, hence on which isoform expression is lost in the brain [11, 13, 14, 17, 18]. *In-vivo* studies in mice demonstrated that full-length *DMD* isoforms (*dp427*) are critical players in GABAergic transmission by contributing to postsynaptic gamma-aminobutyric acid A receptor (GABAARs) clustering [19].

Their deficiency in the *mdx23* mouse model, which lacks *dp427*, was associated with longitudinal changes in brain morphology, resulting in progressive cognitive impairments [20–23]. Similarly, a significant reduction of cortical GABA_A_ receptors in DMD patients, as compared to age-matched normal subjects, was observed and consistent with prefrontal cortex dysfunction [24].

GABAergic and glutamatergic transmission as well as behavioural and cognitive phenotypes were shown to be impaired in the *mdx52* mouse model, deficient in both *dp140* and *dp427* [25, 26]. In accordance with the mouse phenotype, DMD patients lacking *dp427* and *dp140* were reported to present reduced cognitive function, high internalising and externalising symptoms, and increased conditioned startle response [11, 12] alongside a reduction in the grey matter and total brain volume detected by MRI [27, 28]. This reduction was greater in patients lacking both *dp427* and *dp140* than *dp427* only [27]. Finally the lack of *dp71* clear contributes to the severity of the neural phenotype in human studies, with the mean IQ of 55 in *dp71* deficient patients [29]. A behaviorual phenotype also characterises the *dp71*-null mouse model, where a selective increase in excitatory transmission at glutamatergic synapses on Purkinje neurons was observed [30].

Despite a clear correlation between deficiency of different isoforms and resulting phenotypes in the human and mouse models [31], there remain significant knowledge gaps regarding the etiopathogenesis underlying CNS co-morbidities in DMD. Current information on expression of DMD isoforms in the human brain is limited to the mining of transcriptomic data from publicly available databases and key limitations include lack of temporal resolution, absence of isoform and exon-specific data [10, 32, 33]. In order to understand the role of DMD in behaviour and cognition, and whether its expression is required for normal development, it will be crucial to generate a comprehensive map of the regional and cellular distribution of dystrophin isoforms in developing and adult human brains. It will also be important to establish whether human neural cells express single or multiple dystrophin isoforms.

Hence our aim was to systematically assess the expression and localisation of the *DMD* isoforms during development and in the adult human brain. We firstly used quantitative reverse transcription polymerase chain reaction (RT-qPCR) across multiple brain regions. We then employed *5’/3’* and isoform-specific *DMD in situ* hybridisation (ISH) probes to map *DMD* transcripts at the regional, cellular and sub-cellular levels. Using multiplex ISH probes, we studied the co-expression of *DMD* isoforms and their co-localisation with excitatory and inhibitory neurons. Specific antibodies were used to map dystrophin protein expression. Our studies provide the first comprehensive spatial and temporal expression patterns of the dystrophin isoforms of the developing and adult, pathology-free human brain.

## Materials and methods

### Human tissues

Human tissue procedures were conducted in accordance with the regulations with informed consent in accordance with the UK Human Tissue Act 2006 for study participation under ethical approval (NRES Committee London—Fulham, UK; REC: 18/LO/0822). The human embryonic and foetal material was provided by the Joint MRC/Wellcome Trust (Grants MR/006237/1, MR/X008304/1 and 226202/Z/22/Z) Human Developmental Biology Resource (https://www.hdbr.org).

Tissue bank provided brain tissue from several donors at 17 (41 days) and 23 (56 days) Carnegie stages (CS) and 10, 14, 15, 19, and 20 post-conception weeks (pcw). Details of the subjects and samples cohorts are indicated in **Supplementary Tab. 1**.

Adult human brains: FFPE and frozen post-mortem brain specimens from four ‘non-neuromuscular’ pathology-free subjects (in **Supplementary Tab. 1**) were selected and procured from the Edinburgh Brain Bank (Ethic Ref 20/PR/0582). All subjects showed no significant abnormalities after histopathological examination of the brain. Tissue samples were obtained from the Edinburgh Brain Bank (EBB) as part of BRAIN UK, which is supported by Brain Tumour Research and has been established with the support of the British Neuropathological Society and the Medical Research Council [34].

Three distinct samples cohorts were available:1) frozen and FFPE specimens from embryonic and foetal subjects; 2) five ‘matching’ brain areas from the same adult subjects available both frozen and FFPE and; 3) twelve remaining adult brain areas available FFPE only.

### Transcript analysis

#### RNA isolation

According to the manufacturer’s instructions, RNeasy Mini Kit (Qiagen, Cat. No. 74104) and RNeasy FFPE Kit (Qiagen, Cat. No. 73504) were used to isolate RNA from frozen samples.

Total RNA concentration was measured using the NanoDrop 2000 spectrophotometer (Thermo Scientific, Cat. No. ND-2000). cDNA was generated by reverse transcription from 200 ng (frozen) and 100 ng (FFPE) of RNA using the High-Capacity RNA-to-cDNA™ Kit (Thermofisher, Cat. No. 4387406) according to the manufacturer’s instructions. FFPE-derived cDNA from adults was pre-amplified with the TaqMan™ PreAmp Master Mix (Thermofisher, Cat. No. 4391128) according to the manufacturer’s instructions.

#### Gene expression analysis by RT-qPCR

Subsequently, *DMD* expression was detected by RT-qPCR with the TaqMan™ Fast Advanced Master Mix (Thermofisher, Cat. No. 4444557) and a set of TaqMan isoform-specific probes (Thermofisher Cat.No 4332079).

Relative expression was calculated by 2−ΔCt formula and normalised to the brain-specific *RPL13* housekeeping gene. *RPL13* was selected as a housekeeping gene following a comparison with other *CYC1*, *GAPDH*, and *UBE2D2* genes (data not showed), demonstrating its higher consistency of expression in human brain samples, in line with previous studies. Data were then converted to a common scale, log2 was transformed and visualised on heatmaps with the following colour code: white= unprocessed; light green=low expression; green=moderate expression; orange/red=high expression; grey=undetermined.

To mitigate the RNA fragmentation-related effects resulting from tissue fixation [35], a combined polyA/random hexamers reverse transcription and a cDNA preamplification steps were introduced for maximising the RT-qPCR sensitivity in FFPE samples [36, 37]. Despite our efforts to design probes and primers amplifying short amplicons (between 21 and 31bp) aimed at making highly sensitive RT-qPCR assays, a longer amplicon length was required for *dp427m* (41bp), *dp427p1* (37bp) and *dp40* (92bp) amplification. Consequently, when FFPE and frozen RT-qPCR results from the same area (and same subject) were compared, we detected a reduction in log2 FC magnitude of *dp427m*, *dp427p1* in the FFPEs areas. This was likely due to the larger amplicon sizes compared to the other isoforms and in particular the full-length *dp427c* (21bp). The impact of RNA fragmentation was more evident with the 92bp amplicon (more than 3x higher than the average) of *dp40* which was detected in frozen samples only. This is in line with previous reports showing that smaller amplicons produce more comparable cycle threshold (Ct) between FFPEs and frozen samples because less prone to fragmentation [36–38].

### *In situ* Hybridisation

#### RNAscope assays

The RNAscope *in situ* hybridisation (ISH) assay with its high specificity and sensitivity allows for the resolution of single mRNA transcripts which typically appear as discrete punctate/particulate foci with a diameter of 0.1-4.3 microns within the cell cytoplasm and/or nuclear domains (**Fig. 3b**). Each dot/particle represents a single mRNA transcript.

We assessed the prevalence and spatial distribution of *DMD* transcripts within cellular subtypes across different regions in the developing and adult human brain. Specifically, we employed a *DMD 5’* probe targeting region (E2-E10) recognising all nascent and mature full-length transcripts (*dp427m*/*dp427c*/*dp427p*) and a *DMD 3’* probe targeting region (E64-E75) recognising all mature dystrophin transcripts (*dp427*, *dp260*, *dp140*, *dp116*, *dp71*/*dp40*) except *dp40* (**Fig. 3a**). Appropriate positive and negative controls were employed in parallel to confirm RNA preservation and non-specific labeling (**Fig. S1b**). Brain FFPE and frozen samples were pre-treated according to the existing manufacturer’s instructions and incubated with probes designed by ACD Bio (Advanced Cell Diagnostics Inc., Newark, CA).

Singleplex RNAscope assays (Advanced Cell Diagnostics Inc, Cat. No. 322360) relying on a custom-made RNAscope *DMD 5’* probe and a commercially available *DMD 3’* probe was performed according to the manufacturer’s protocol. Probes are listed in **Supplementary Tab. 2**.

Before being processed for *in situ* assays, selected control sections were incubated against a commercially available RNAscope *UBC* (ubiquitin C) probe to confirm RNA preservation and PBS/probe diluent to test the extent of the background signal (negative control). Tissue staining was performed according to the manufacturer’s instructions.

#### Basescope assays

Singleplex Basescope assays (Advanced Cell Diagnostics Inc., Cat. No. 323900) combined with a set of singleplex probes were employed to deconvolute the RNAscope signal and study the spatial expression pattern of *dp427m*, *dp427c*, *dp427p2*, *dp140*, *dp71*, and *dp40* in selected brain areas. Tests were performed according to the manufacturer’s protocol (Advanced Cell Diagnostics Inc., Cat. No. 323900-USM). Probes are listed in **Supplementary Tab. 2**. A few steps were modified for embryonic/foetal tissue to avoid tissue digestion. Specifically, the target retrieval was performed for 8 mins in boiling water; tissue was treated with Protease III for 30 min at 40° in high-humidity conditions. Before being processed for *in situ* assays, selected control sections were incubated against a commercially available a Basescope *UBC* (ubiquitin C) probe to confirm RNA preservation and PBS/probe diluent to test the extent of the background signal (negative control). Tissue staining was performed according to the manufacturer’s instructions.

### Tissue imaging and analysis

Whole section images were acquired within 24h at 40x magnification on a NanoZoomer S60 slide scanner (Hamamtsu, Cat. No. C13210-01).

RNAscope: given the large number of sections to be evaluated and complexity of the histological elements in different brain areas, the RNAscope signal was analysed by the neuropathologist only, who scored all the different areas based on the presence of the signal and data were plotted on a heatmap showing the following colour code: white: unprocessed; light green=sporadic (0-5% positive cells); green=low expression (5-20 % positive cells); intense green=moderate expression (20-50 % positive cells); orange=frequent (50-70% positive cells); red=strong expression (>70% positive cells); grey: undetermined.

Singleplex basescope signal semi-quantification: anatomical regions for the processed brain area were manually annotated by a neuropathologist. For each annotated region, a set of images (from three randomly selected fields of view) was exported, and the *in situ* signal was manually scored by three independent operators as indicated in the manufacturer guidelines. We then averaged the results obtained by the three operators and plotted the data into heatmaps. Specifically, we quantified the number of cells displaying at least one dot (positive) and calculated the P+ parameter, representing the percentage of positive cells out of the total cells. Heatmap colour code: white=unprocessed; light green=low expression; green=medium expression; orange/red=high expression; grey=undetermined. Duplex Basescope was performed in a qualitative fashion to detect the co-expression of multiple *DMD* cellular transcripts by probes targeting the following isoforms: *dp427m* (channel 1-green), *dp427c* (channel 2-red), *dp140* (channel 1-green and channel 2-red), *dp71* (channel 2-red), *UBC* (channel 1-green), and *PPIB* (channel 2-red).

### Protein analysis

#### Protein extraction

Protein samples were prepared on ice for extraction after adding 10 μL of extraction buffer (50 μL of protease and phosphate inhibitor mini tab (Pierce, Cat. No. A32959) to 950uL of T-PER tissue protein extraction reagent (Thermo Scientific, Cat. No. 78510) for each mg of tissue. Tissue was then homogenised using Fisher brand 850 Homogeniser (Fisher scientific, Cat. No.15-340-169) for 20 seconds at 10,000 (brain samples) and placed in an ultrasonic ice bath (Branson Ultrasonics, Cat. No. 2510-DTH) for 5 minutes. The homogenates were then centrifuged for 15 minutes at 14,000 g at 4°C, the supernatant was then centrifuged again at 14,000 g at 4°C for 5 minutes and kept at –70°C. Protein extracted from the samples were quantified using RC DC Protein Assay Reagents Package (Bio-Rad Laboratories Ltd, Cat. No. 5000120,).

#### Capillary Western Immunoassay (Wes)

Simple western blot (WES, Bio-techne) was used to quantify Dp427, Dp140 and Dp71 levels in protein lysates from brain samples. Ab154168 (abcam) was used as the antibody with a concentration of 1:1500, which recognises Dp427, Dp260 and Dp140. A total of 2 μg of protein for each sample was prepared for WES analysis following protocols for the 66-440 kDa separation module (Bio-techne, Cat. No. SM-W005) and for the total protein detection module (Bio-techne, Cat. No. DM-TP01). Each dystrophin protein signal was normalised to the total protein detection signal.

#### Protein localisation by immunohistochemistry (IHC)

Tissue was obtained from HDBR or Brain tissue bank as formalin-fixed paraffin-embedded (FFPE) blocks or 5 µm thick paraffin sections on microscope slides. After cutting, slides were placed in an oven or hotplate for 10 minutes. Then, they were loaded onto the Ventana Discovery Ultra machine. First, three cycles of deparaffinisation were performed for 4 minutes each using EZ Prep at 65°C. Heat Induced Epitope Retrieval (HIER) is performed using Citrate-based buffer pH 6.0 at 95°C for three cycles (8min, 12min, 20min). Slides were washed three times using a reaction buffer (Tris-based containing ∼2% Brij 35 and ∼4% Acetic Acid). An amplification kit was used for primary anti-dystrophin antibodies. Two primary anti-mouse antibodies were used, one toward the C-terminus (1:150), detecting all dystrophin isoforms except dp40 (Leica Biosystems, NCL-DYS2), and the second against the rod domain (1:150) detecting longer dystrophin isoforms (NCL-DYSA, Leica Biosystems). The primary antibodies incubated for 60 minutes involved heating at RT. A secondary antibody was performed using OmniMap anti-Mouse HRP (Roche diagnostics, Cat. No. 760-4310), and then a Chromomere DAB kit (Biocare Medical Cat. No. ACT500) was used for nuclear staining. Finally, sections were mounted after three times washing for 2 minutes each and left to dry overnight.

### Statistical analysis

Significant changes in transcript levels in brain tissue for dystrophin expression were assessed by one-way analysis of variance (ANOVA) to compare the differences between multiple groups (**Supplementary Tab. 3**). Statistical analysis was performed using the Prism 9 software (GraphPad Software LLC, Cat. No. 140000). Data are presented as mean ± SEM or as a heatmap. The p-value < 0.05 was reported as significant ** p≤0.01, *** p≤0.001 and **** p≤0.0001.

## Results

### Dystrophin isoform expression in human brains from embryo to adult

To gain insights into the regulation of dystrophin isoforms in developing and adult human brains, we first assessed their expression by RT-qPCR. Specifically, we quantified regional expression of *dp427c*, *dp427m*, *dp427p1*, *dp427p2*, *dp260-1*, *dp260-2*, *dp140*, *dp116*, *dp71*/*40* (sharing the first unique exon) and *dp40* isoforms in embryonic, foetal, and adult frozen and/or formalin-fixed and paraffin embedded (FFPE) post-mortem pathology-free brain samples. RT-qPCR data showing isoform brain expression from embryo to adult are summarised in the compact heatmap in **Fig. 2** and **S1a**.

**Fig. 2.**
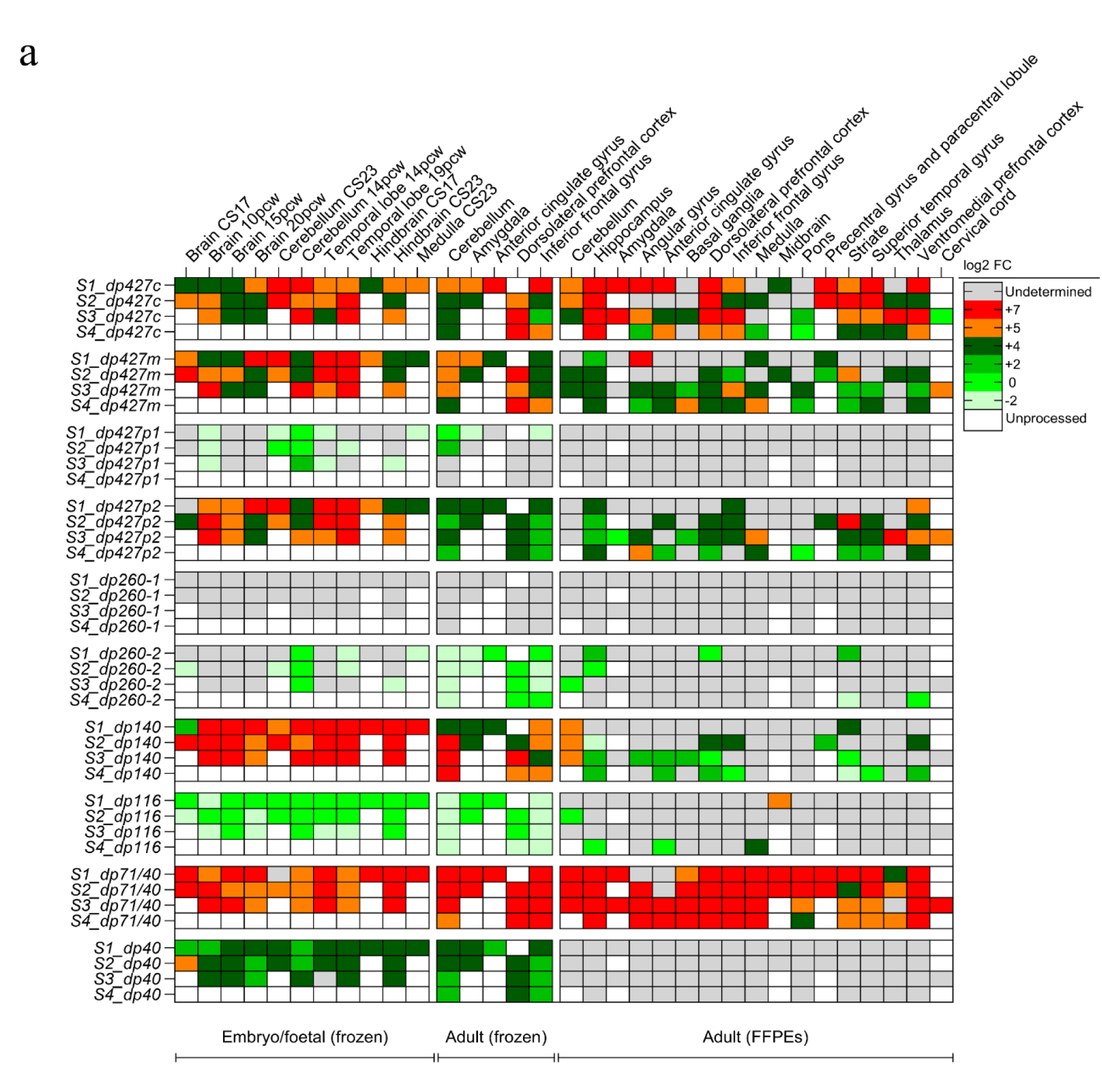
Expression patterns of the main *DMD* isoforms in developing and adult brain areas assessed by RT-qPCR. Heatmap showing the log2 fold changes of the *DMD* isoforms in prenatal and adult human brains. Gene expression was normalised to the brain specific *RPL13* housekeeping gene. Colour code: white: unprocessed; light green: low expression; green: moderate expression; orange/red: high expression; grey: undetermined. Pcw: postconceptional weeks; CS: Carnegie stage. FFPEs: Formalin Fixed Paraffin Embedded tissue samples; S1-4: subject. Log2 fold changes values are shown as mean ± standard error.

Among the full-length isoforms, the Purkinje *dp427p1* transcripts were hardly detectable or absent in developing and adult brains. *Dp427p2* showed high expression in the embryonic and foetal brain, including cerebellar and extra-cerebellar regions, but was significantly downregulated in the adult brain. The cortical *dp427c* isoform was highly expressed in the developing brain from the earliest embryonic stage studied (CS17) and its expression was maintained in most areas of adult brain, with the highest levels observed in hippocampus, amygdala, and several neocortical areas including the prefrontal cortex. Moderate to high levels of expression were also observed in the cerebellum. *Dp427m* transcripts were highly expressed throughout development and showed variable expression levels (low/moderate to high) in the adult brain.

The d*p260-1* splicing isoform was undetectable at all stages, whereas a low expression of *dp260-2* was detected in some developing and adult brain regions.

Among the intermediate isoforms, *dp140* was highly expressed in the embryonic and early foetal brains without regional specification, but not in the adult brain, where it became largely restricted to the cerebellum with high expression levels; lower levels of expression were detected in the dorsolateral prefrontal cortex, striate cortex and inferior frontal gyrus. Low *dp116* expression was detected in most of the frozen regions analysed.

Both short isoforms *dp71/dp40*, showed high expression levels in the embryonic and early foetal brains, and continued to remain high in the adult brain, with little regional specification. When the 5′UTR-*dp71* and the 3′UTR-*dp40* were targeted to specifically assess *dp40* transcripts, low *dp40* expression was detected in both embryonic/foetal and adult frozen specimens, but not in adult fixed tissues (**Fig. S1a**), where higher RNA fragmentation can occur as explained in Materials and methods.

In summary, all the *DMD* isoforms apart from *dp427p1* and *dp260-1* are consistently expressed in the human brain, with changes in *dp427p2* and *dp140* expression suggesting developmental regulation of these isoforms.

### RNAscope *in situ* hybridisation assay to *5’* and *3’* regions of *DMD* mRNA allows spatial mapping of dystrophin transcripts in the developing and adult human brain

Labeling with the *DMD 5’* probe, targeting the full-length isoforms, showed a consistent pattern across all neuronal populations in the form of a discrete, large intranuclear particulate signal measuring up to 4.3 µm, suggestive of transcriptional ‘hot spots’ containing multiple nascent transcripts (**Fig. 3b)**. A few neurons also displayed smaller, punctate foci in the nucleus and/or cytoplasm. Labeling with the *DMD 3’* probe, targeting all the isoforms apart from *dp40*, showed 1-3 large discrete intranuclear foci, and multiple smaller, mostly cytoplasmic foci, suggestive of mature transcripts (**Fig. 3b)**. **Fig. 3c** shows representative images and a heatmap summarising the semi-quantitative values from the RNAscope signal scored by the pathologist in the adult brain, as detailed in the Materials and methods section.

**Fig. 3.**
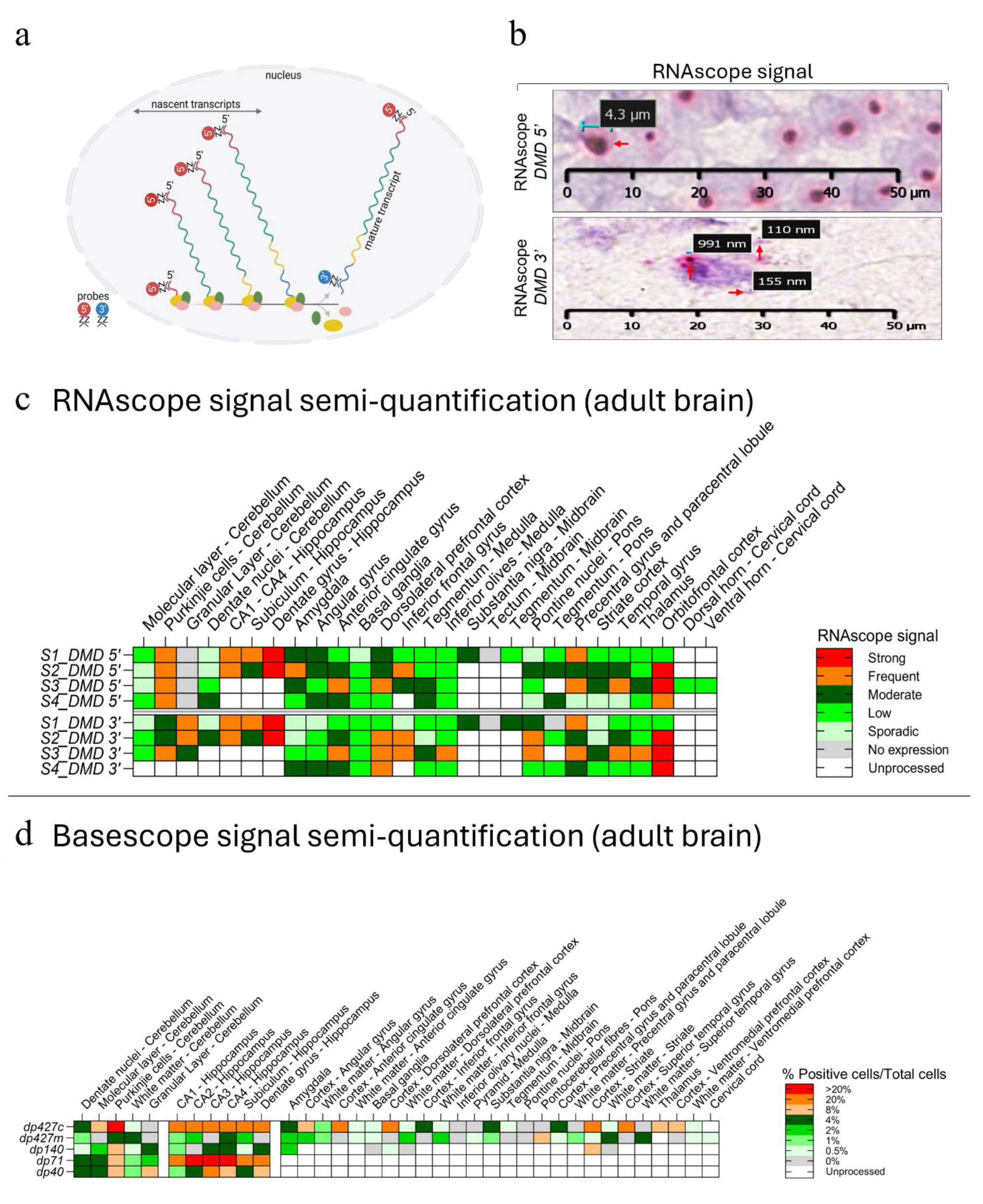
*In situ* hybridisation in the human brain. RNA labelling dynamics at the *5’* and *3’* ends of the *DMD* gene. The *DMD 5’* probe detects *dp427c*, *dp427m* and *dp427p* and the *DMD 3’* probe detects all the mature *DMD* transcripts except *dp40* (**a**). Images show the variable size of the RNAscope red punctate dots detected using *DMD 5’* and *DMD 3’* probes (**b**). Heatmap representing the *DMD 5’* and *DMD 3’* RNAscope signal scores in each adult ROIs (**c**). Heatmap representing the singleplex Basescope P+ (% Positive cells/total cells) across different adult brain areas (**d**). Colour code: white: unprocessed; light green: low expression; green: moderate expression; orange/red: high expression; grey: undetermined; S1-4: subject.

In CS17 embryos, more extensive labeling was noted with the *DMD 3’* probe than the *DMD 5’* probe indicative of higher expression of the shorter-length *DMD* isoforms (**Fig. 4** and **Fig. S2a**). The *DMD 3’* signal was abundant in the ventricular and subventricular zones of the telencephalon (**Fig. S2a**), the cerebellum (**Fig. 4a** and **S4a**), pons (**Fig. 4b**) and medulla (**Fig. 4d**), spinal cord and dorsal root ganglia, suggestive of *DMD* expression in the neural progenitor cells and early migratory neurons. *DMD 5’* signal was localised to the pontine (**Fig. 4b**) and medullary neuroepithelial cells (**Fig. 4d**), possibly reflecting the expression of full-length isoforms. The optic vesicle exclusively expressed the *DMD 3’* signal at CS17 (**Fig. 4f**), whereas, the trigeminal ganglion (**Fig. 4h**) and skeletal muscles (**Fig. 4g**) expressed both *DMD 5’* and *DMD 3’* labeled transcripts.

**Fig. 4.**
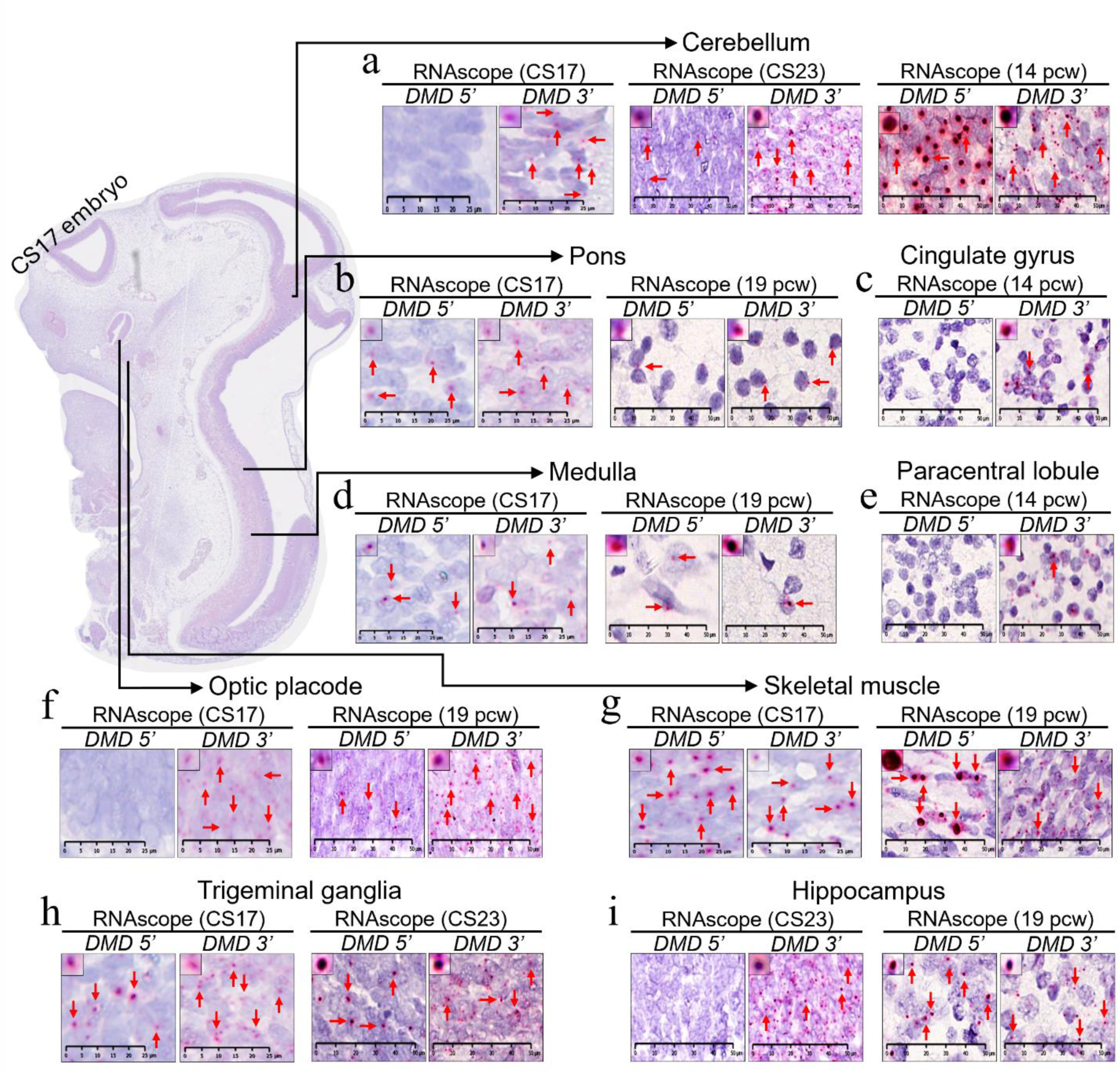
RNAscope signal in the embryonic and foetal human brain. Representative images showing stable RNAscope signal in both embryonic and foetal FFPE brain samples (**a**-**i**). The top left panel shows the most representative puncta to a higher magnification. Each red arrow indicates RNAscope signal (red dots) or representative dots in case of high RNA expression/clusters. Pcw: postconceptional weeks; CS: Carnegie stage; Hippocampus CA: Cornu Ammonis. Scale bars = 25 and 50 µm.

At the late embryonic stage, CS23, *DMD 3’* labeled transcripts continued to be widely distributed in cerebellum (**Fig. 4a**, **S2b** and **S4a**), hippocampus (**Fig. 4i** and **S2b**), amygdala, cerebral cortex, basal ganglia, medulla and retina (**Fig. S2b**). As at CS17, skeletal muscle expressed both full-length and shorter transcripts (**Fig. 4g** and **S2b**). Overall, the expression of *DMD 5’* labeled full-length transcripts increased with development and was already apparent at CS23 in all the above regions studied except the hippocampus **(Fig. 4i)**.

At foetal stages (pcw 14 and pcw 19), *DMD 5’* labeled transcripts were also detected in the hippocampus (**Fig. 4i** and **S2d**), in addition to cerebral cortex (**Fig. S2d**). The *DMD 5’* signal was particularly striking in the Purkinje cells and in deep nuclei of the foetal cerebellum where large puncta were observed (**Fig. 4a**, **S4a** and **S2c**).

In the adult brain *DMD 5’* and *DMD 3’* labeled transcripts were widely expressed across several neocortical regions: primary motor cortex, dorsolateral and ventromedial prefrontal cortex, inferior frontal gyrus, superior temporal gyrus, angular gyrus, calcarine/striate cortex and anterior cingulate gyrus (**Fig. 5d, 5f, 5g** and **S3a**). The expression pattern was similar across these regions with pan-cortical localisation of transcripts. Within the cortices, most transcripts were neuronal, including majority of pyramidal cells (**Fig. 5** and **S3a**) and a small proportion of the *5’* transcripts localised to oligodendroglia-like cells in the cortex and white matter (**Fig. S3a**). Both *DMD 5’* and *DMD 3’* labeled transcripts were localised to the upper motor neurons, the giant pyramidal cells of Betz **(Fig. 8e)**.

**Fig. 5.**
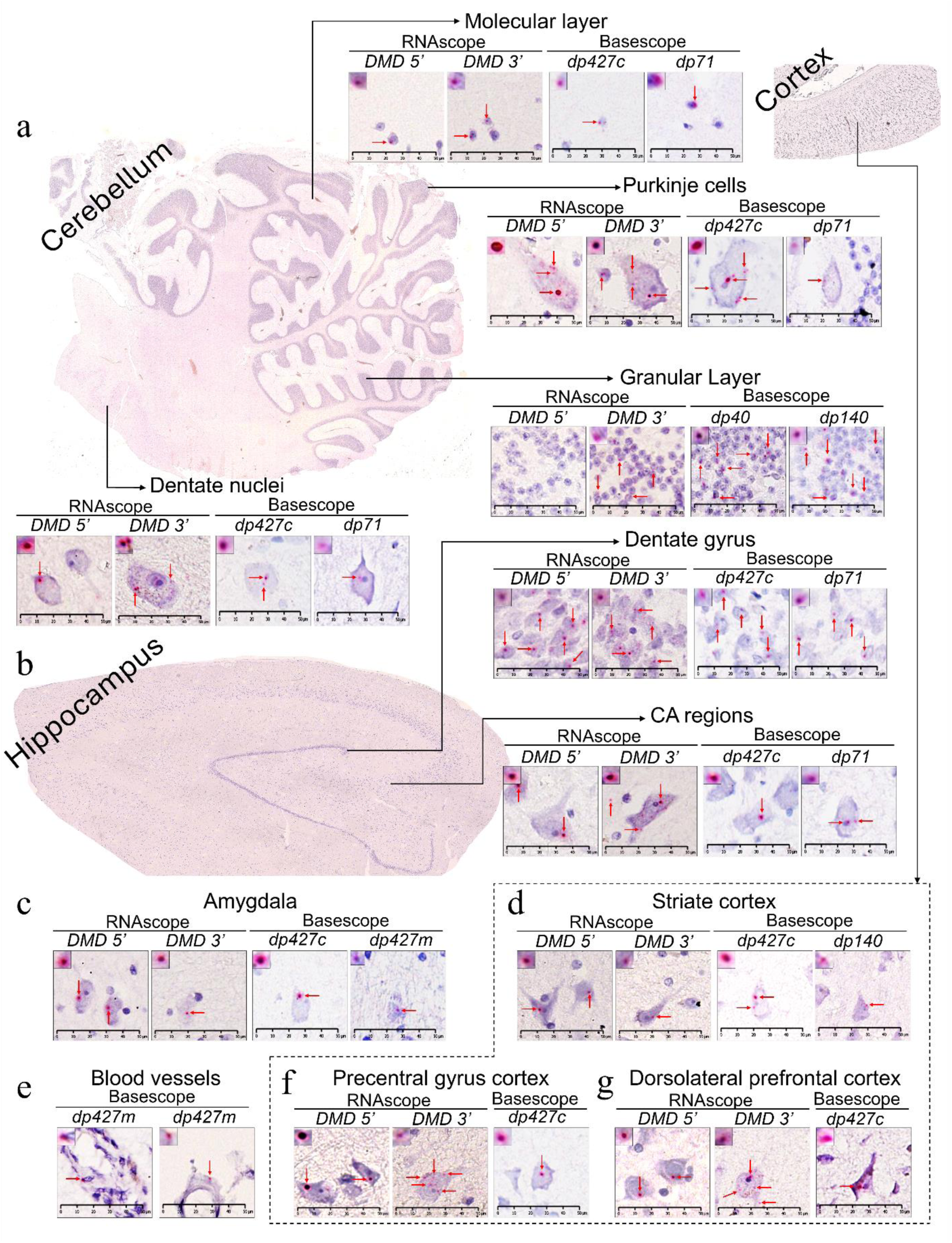
RNAscope and singleplex Basescope signal in the adult human brain. Representative images of adult brain areas hybridised with either *DMD 5’* or *DMD 3’* probes or singleplex Basescope probes (counterstained with haematoxylin) showing stable RNAscope signal in FFPE brain samples (**a**-**g**). Each red arrow indicates RNAscope/Basescope signal (red dots) or representative dots in case of high RNA expression/clusters. The top left panel shows the most representative puncta to a higher magnification. CA: Cornu Ammonis. Scale bars = 50 µm.

*DMD* transcripts were abundant in the hippocampus proper CA1-CA3, CA4/hilar region and the dentate fascia **(Fig. 5b and S3a)**. In the cerebellar cortex, *DMD 5’* and *DMD 3’* labeled transcripts were noted in the molecular and Purkinje cell layers **(Fig. 5a, S3a** and **S4b)**, whereas only *DMD 3’* labeled transcripts were seen in the granule cell layer **(Fig. 5a, S3a** and **S4b)**, suggesting exclusive expression of shorter mature transcripts in these cells (*dp140*/*dp71*). Both *DMD 5’* and *DMD 3’* labeled transcript were seen in the dentate nucleus **(Fig. 5a)**.

While both *DMD* 5’ and 3’ transcripts were localised within the amygdala, subregional (nucleus-wise) specification could not be ascertained due to suboptimal orientation of the blocks. High expression was detected in the prefrontal cortices, inferior frontal gyrus, hippocampus and the cerebellum **(Fig. 3c**).

### Basescope *DMD* isoform-specific, singleplex and multiplex *in situ* hybridisation assays reveal isoform diversity and coexpression within neuronal, glial and vascular domains across key brain regions

*Singleplex Basescope*:While the analysis with RNAscope *DMD 5’* and *DMD 3’* probes could demonstrate differences in localisation of dystrophin long and short isoforms, it did not provide information on expression and localisation of specific *DMD* isoforms. Hence, we used singleplex Basescope assays to localise dystrophin isoforms, *dp427c*, *dp427m*, *dp140*, *dp71*, and *dp40.* We could not study *dp427p2* due to lack of specificity of the labelling probe. Expression of these isoforms was semi-quantified in adult brains and shown as a heatmap (**Fig. 3d**), and examples of cellular localization are shown in **Fig. 5** and **S3b**. In the adult brain, *dp427c* transcript was widely expressed across all neocortical regions (**Fig 5d, 5f** and **5g**), and also in the hippocampus (**Fig. 5b**), amygdala (**Fig. 5c**) and cerebellum (**Fig. 5a**). *Dp427m* was consistently expressed within the leptomeningeal and cortical blood vessels (**Fig. 5e and S3b**) containing a muscular coat (most likely in the smooth muscle cells).

*Dp140* transcripts were expressed in the dorsolateral and ventromedial prefrontal cortices and the striate cortex (**Fig. S3b**). *Dp71* was detected in the ventromedial orbitofrontal cortex with transcripts identified in pyramidal cells.

In the adult hippocampus, the granule cells in the dentate gyrus showed moderate to frequent expression of *dp427c* (**Fig. 5b** and **S3b**), *dp140* (**Fig. 6b**) and *dp71* (**Fig. 5b** and **S3b**) and low levels of *dp427m* and *dp40* transcripts. The hilum/CA4, CA1-CA3 and subiculum showed moderate expression of *dp427c* and *dp71* (**Fig. 5b** and **S3b**), and low level of *dp427m* transcripts (**Fig. 3d**).

**Fig. 6.**
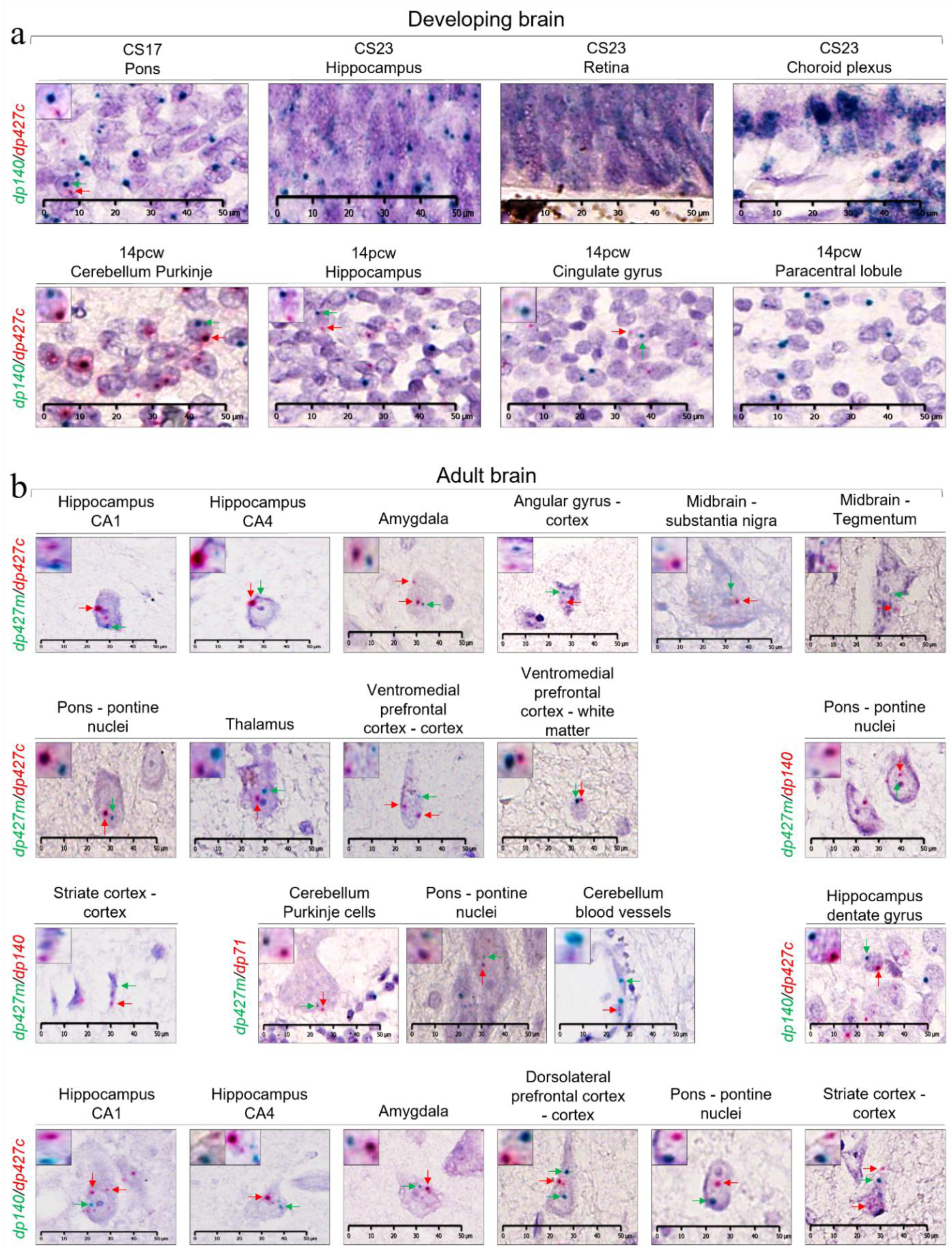
Detection of co-expressed *DMD* isoforms in developing and adult human brain areas by duplex Basescope. Representative images of developing brain samples hybridised with the *dp140* (green) and *dp427c* (red) probes. Note some cells express both isoforms (**a**). Representative images of adult brain samples hybridised with four different probe combinations: *dp427m*(green)/*dp427c*(red); *dp427m*(green)/*dp140*(red); *dp427m* (green)/*dp71*(red); *dp140* (green)/*dp427c*(red) (**b**). Red and green arrows indicate RNA transcripts (dots) co-expressed in the same cell. The top left inset shows the co-expression events at a higher magnification. bv: blood vessels; Pcw: postconceptional weeks; CS: Carnegie stage; Hippocampus CA: Cornu Ammonis. Scale bars = 50 µm.

The amygdala showed low levels of *dp427c* and *dp140* expression (**Fig. 5c** and **6b**). Very few neurons expressed *dp427m* (**Fig. 5c**). Sparse *dp427c* transcripts were noted in the basal ganglia. Few oligodendroglial cells expressed *dp427m* in the white matter (**Fig. S3b)**. The thalamus showed moderate expression of *dp427c* (**Fig. S3b**). *Dp427m* signal was noted mostly in oligodendroglia, and rarely in neurons.

In the midbrain, dystrophin transcripts were mainly present in the substantia nigra pigmented neurons, specifically, *dp427c,* with occasional neurons expressing *dp427m*. The dorsal midbrain and the midbrain tegmentum showed occasional *dp427c* and *dp427m* in the Edinger Westphal nuclei (**Fig. S3b**), oculomotor nerve nuclei and the mid-line raphe neurons.

The pontine tegmentum, including the pigmented neurons of the locus coeruli, the periventricular grey, the midline raphe nuclei and the pontine reticular formation showed modest expression of *dp427c* and *dp140* transcripts.Similarly in the basis pontis, there was moderate expression of *dp427c*, *dp427m, dp140* and *dp71* transcripts in the pontine nuclei. *Dp427m* expression was also noted in the oligodendroglial cells within the white matter tracts (corticopontine, corticospinal and pontocerebellar fibres) (**Fig. S3b**).

In the medulla, sparse *dp427c* transcripts were localised to the medullary tegmentum (hypoglossal nuclei and reticular formation) and the inferior olivary nucleus. *Dp427m* was expressed in oligodendroglial cells in the pyramidal tracts and the emerging rootlets of the vagus nerve.

The adult cerebellum showed few *dp40* and *dp71* (**Fig. 5a**) transcripts expressed within the molecular layer (likely in interneurons), Purkinje cells, dentate nucleus (**Fig. 5a**) and granule cells. *Dp71* transcripts were also seen within the parenchymal capillary walls and the pia-glial membrane (glia limitans) (**Fig. 9h** and **9i**), but not within the leptomeningeal blood vessels. In the cerebellum, *dp140* transcripts were detected in a moderate number of granule cell layer (**Fig. 5a**). A small number of molecular layer neurons, Purkinje cells and dentate neurons also expressed *dp140* transcripts. Moderate numbers of Purkinje cells and neurons in the dentate nucleus and few interneurons in the molecular layer expressed *dp427c* (**Fig. 5a**). In contrast, there was no *dp427c* signal in the granular cell layer. *Dp427m* expression was restricted to sparse basket cells within the Purkinje cell layer. **(Fig. S5a)**.

*Duplex Basescope*: In addition, duplex Basescope with *dp427m*/*dp427c*, *dp140*/*dp427c*, *dp427m*/*dp140* and *dp427m*/*dp71* probes was used to further assess isoform localisation in the adult brain, and establish whether more than one dystrophin isoform may be expressed in the same cell (**Fig. 6** and **S5**).

In the adult brain, neuronal *dp140/dp427c* co-expression was seen in the, hippocampus, pons, striate cortex and dorsolateral prefrontal cortex while rare neuronal events were noted in pyramidal cells across all hippocampal subsectors and in the amygdala (**Fig. 6b**). Co-expression of the two full length *dp427c*/*dp427m* isoforms was seen in one of the midbrain’s raphe neurons (**Fig. 6b**). Unexpectedly, moderate *dp427m* signal was noted in oligodendrocytes, both within the cerebellar white matter, the cortex and the underlying white matter across all neocortical areas (**Fig. 6b** and **S5a**).

Due to the limited number of prenatal sections available, we elected to study CS17, CS23 and 14 pcw sections for duplex BaseScope with the *dp140/dp427c* isoforms as they were the most relevant neuronal transcripts previously detected by RT-qPCR. In these sections, we detected a robust *dp140* expression (**Fig. 6a**), consistent with the RT-qPCR data and in line with previous literature findings, indicating *dp140* abundancy during the early stages of life. At embryonic CS stages 17 and 23 *Dp140* transcripts were most abundant and widely expressed in the developing CNS, including the telencephalic cortical plate, dorsal hippocampus, thalamic, mesencephalic, cerebellar and brain stem neuroepithelia. *Dp140* transcripts were noted in the subventricular germinal neuroepithelium as well as the early migrating neurons. Co-expression of *dp140/dp427c* isoforms was also detected in the CS17 embryonic pons and during the foetal 14pcw stage in cerebellum, hippocampus and cingulate gyrus (**Fig. 6a**). *Dp427c* transcripts were also ubiquitous, but fewer compared to *Dp140*, except in the mesencephalon where they were seen in abundance.

At 14 pcw, the developing cerebellum showed preponderance of *Dp427c* transcripts in the Purkinje cell zone, whereas *Dp140* transcripts were more frequent in the external granule cell layer and the dentate nucleus. Expression of *Dp71* and *Dp40* was not assessed in the embryonic and foetal samples.

Altogether, results from singleplex and duplex Basescope assays confirmed that *dp427c* is abundantly expressed in neurons across all brain regions from early development to adulthood. *Dp427m* is expressed mainly in the leptomeningeal blood vessels, and small number of oligodendroglial cells and neurons. *Dp140* is highly expressed in embryonic and foetal brains and continues to be expressed at low to moderate levels in the adult forebrain, cerebellum and the brain stem. *Dp71* is widely expressed in neurons, as well as the parenchymal microvasculature and the glia limitans. The adult human hippocampus (dentate granule cells), cerebellum (Purkinje cells) and pons (neurons in the pontine nuclei) show the greatest diversity of dystrophin isoform expression.

### *DMD* expression in excitatory and inhibitory neurons

Due to constraints in multiplex immunohistochemistry in FFPE tissues, we used duplex RNAscope comprising dual labeling of *DMD 5’* and *DMD 3’* probes with *GAD1* as a proxy for GABAergic inhibitory neurons and *SLC17A7* as a proxy for glutamatergic excitatory neurons. In the 14 pcw cerebellum, the migrating Purkinje cells coexpressed *GAD1* and *DMD 3*’ transcripts (**Fig. 7a** and **7b**). In contrast, in the granular cell layer, few cells coexpressed *SLC17A7* and *DMD 3’* transcripts (**Fig. 7c** and **7d**). In the adult cerebellum, this distinction was even more pronounced, with coexpression of *DMD 5’* and *GAD1* transcripts in the molecular layer interneurons (**Fig. 7k**), Purkinje cells (**Fig. 7e**) and majority of cells in the dentate nucleus (**Fig. 7f**), whereas *DMD 3’* and *SLC17A7* transcripts were coexpressed mainly in the granule cells (**Fig. 7i**) and a small number of neurons in the dentate nucleus.

**Fig. 7.**
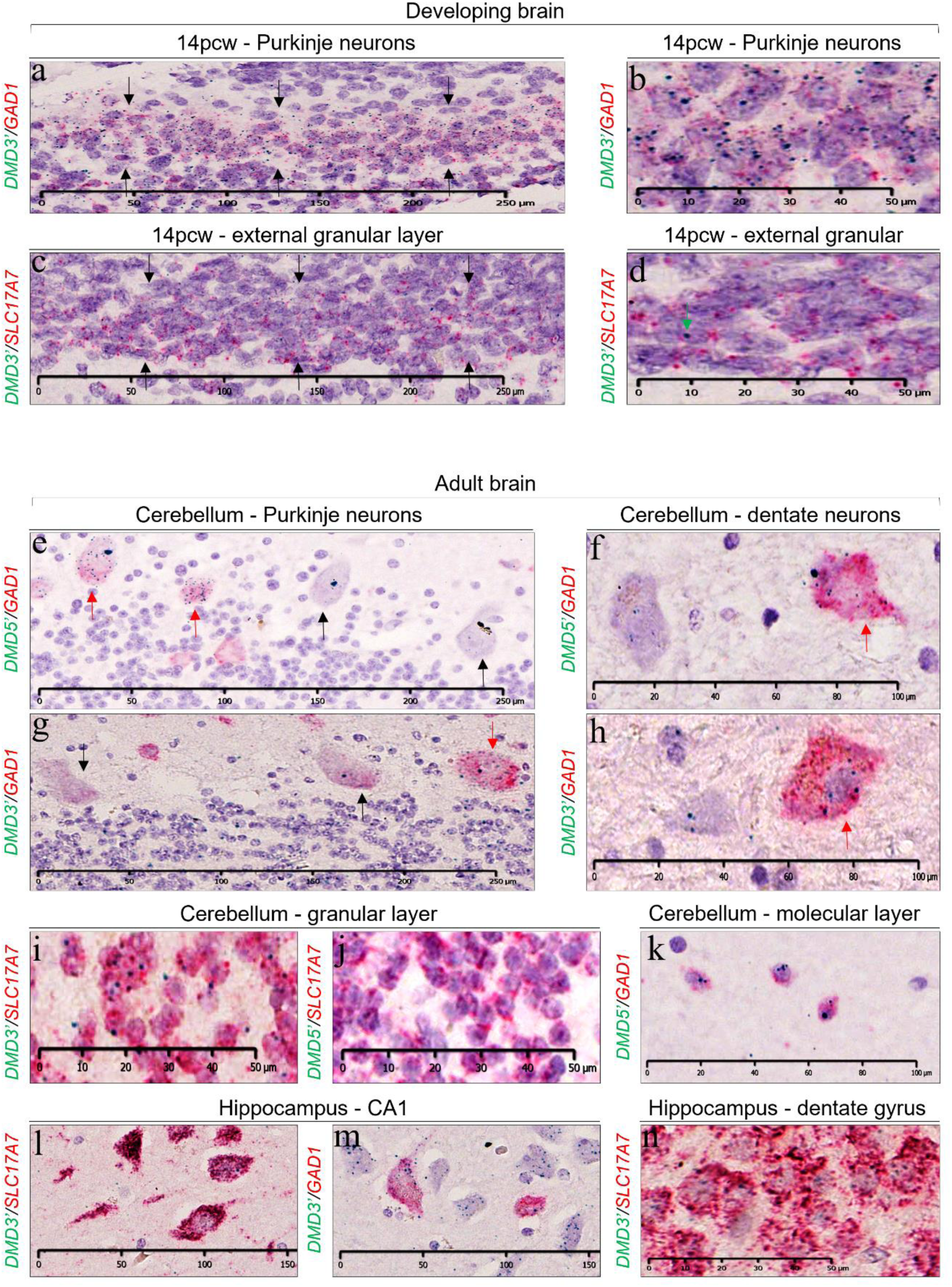
Detection of *DMD* isoforms, GABAergic and glutamatergic neurons in developing and adult human brain areas by duplex RNAscope. Representative image of experiments performed in 14pcw cerebellum (**a**-**d**). Representative images of the results from the adult samples (**e**-**n**). Green dots indicate RNAscope *DMD 5’* and *DMD 3’* transcripts. Red signal indicates *GAD1* (GABAergic) or *SLC17A7* (glutamatergic) transcripts. Black arrows indicate migrating Purkinje neurons (**a**) external granular layer (**c**) *GAD1*- or low *GAD1*+ Purkinje neurons (**e**, **g**). The green arrow shows sporadic *DMD 3’* signal in **d**. Red arrows indicate *GAD1*+ Purkinje neurons (**e**, **g**) and *GAD1*+ dentate neurons (**f**, **h**) in the cerebellum.

In the adult hippocampus, an overwhelming majority of dentate neurons and pyramidal cells (CA1-CA3) coexpressed *DMD3’* and *SLC17A7* transcripts (**Fig. 7l** and **7n**), and a small number of cells, most likely interneurons coexpressed *DMD 3*’ and *GAD1* transcripts (**Fig. 7m**). A similar pattern was evident in the neocortical areas examined, with most pyramidal cells coexpressing *DMD 3’* and *SLC17A7* transcripts (**Fig. S6b-f**). Notably, there was no evidence of labeling in the subcortical and/or deep white matter with any of the probes above.

### Immunohistochemical localisation of the dystrophin protein in the developing and adult human brain

In order to establish whether there was concordance of dystrophin transcript and protein expression, developing and adult brains were immunostained with the antibodies to dystrophin, DYS2, that reacts with the C-terminus, hence detects all isoforms apart from dp40, as well as DYSA. The latter antibody recognises an undisclosed epitope in the rod domain, but previously characterised in mice, and shown to detect both full-length (dp427) and shorter (dp71) isoforms of the dystrophin protein [31].

At stage CS17, DYS2 reactivity was detected in the cerebellum, pons, and medulla, whereas DYSA staining was negative (**Fig. 8a**), indicating that the longer isoforms are not present, or their expression is below the sensitivity of the method, at this early developmental stage. Similarly, only DYS2 reactivity was observed in the CS23 cerebellum (**Fig. 8b**), whereas weak DYSA reactivity was detected in the CS23 trigeminal ganglia (**Fig. 8b**). At the foetal stage examined, 14pcw, both cerebellum and hippocampus showed some DYS2 staining but very weak DYSA reactivity, suggesting predominant expression of shorter isoforms (**Fig. 8c**).

**Fig. 8.**
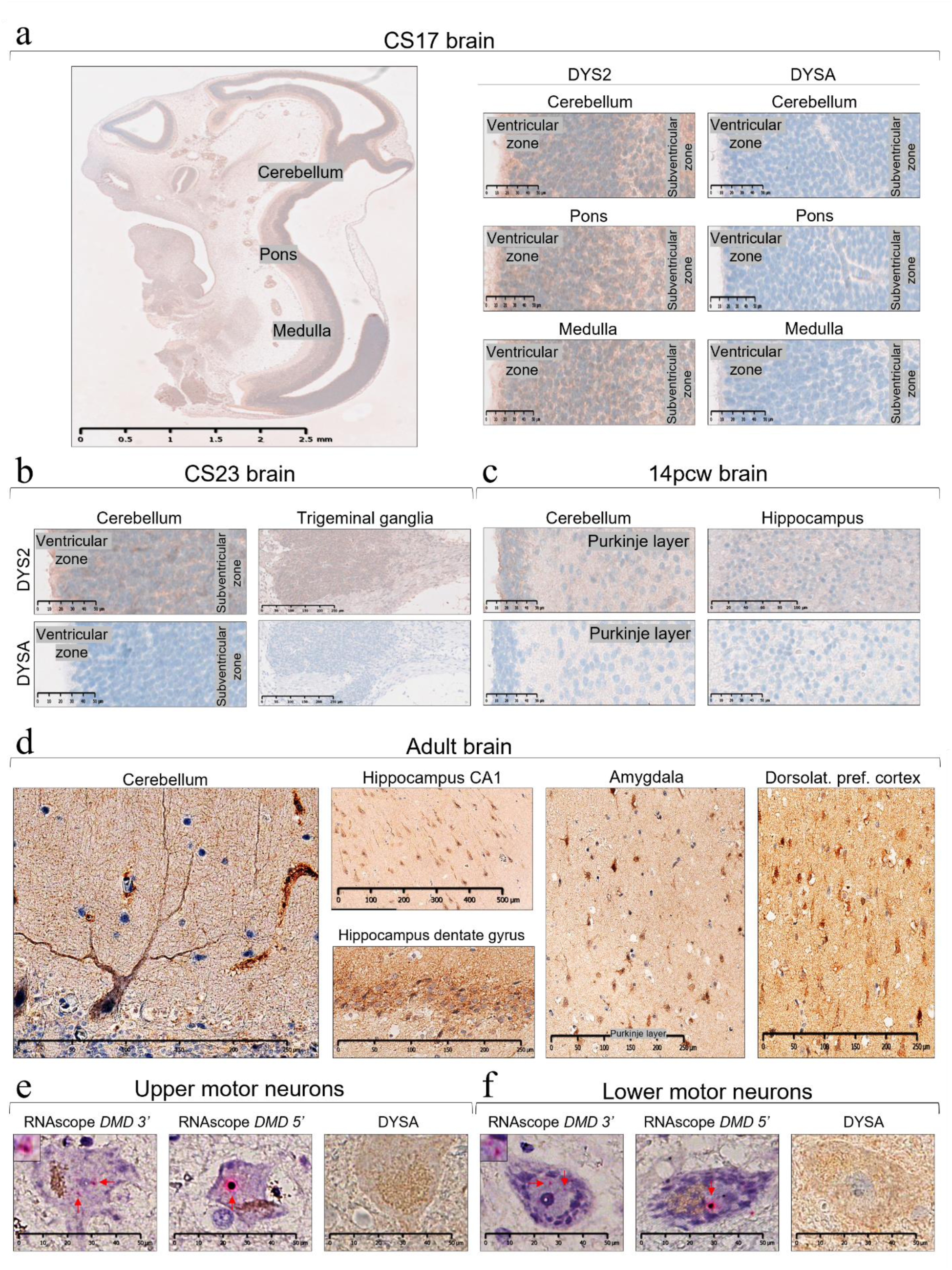
Dystrophin detection by immunohistochemistry in developing and adult brain regions. Developing brains stained either with the monoclonal antibody DYSA (dystrophin N-Terminus) or DYS2 (dystrophin C-Terminus) (**a**-**c**). Adult brain sections stained with DYSA (**d**). Adult upper and lower motor neurons stained with DYSA and RNAscope *DMD 5’* and *DMD 3’* probes (**e**-**f**). Pcw: postconceptional weeks; CS: Carnegie stage.

In contrast, in adult brains DYSA strongly stained several neuronal cell types. Dystrophin expression across all neocortical regions was pan-cortical (**Fig. 8d**), sometimes with accentuated labelling in layer II and layer V/VI. The expression pattern was more readily discernible within the pyramidal cells, partly owing to the size of these neurons. Moderate to strong labelling was observed within the neuronal cell body, with accentuated punctate distribution along the cell surface and the dendritic trees, whereas the axons were negative. There was strong dystrophin expression in the upper motor neurones, so-called giant pyramidal cells of Betz, within the precentral gyrus (**Fig. 8e**).

In the hippocampus, dystrophin immunoreactivity was noted within the dentate granule cells, hilar/CA4 region, Ammon’s horn (CA1-CA3) and the subiculum, restricted to the neuronal somata and their dendritic trees (**Fig. 8d**). Within the amygdaloid complex the presence of large amounts of lipofuscin pigment confounded ascertainment of dystrophin signal within the neurones (**Fig. 8d**).

In the cerebellum, Purkinje cells displayed a distinct punctate labelling pattern at the surface of the cell bodies and their dendritic arbour (**Fig. 8d**). Small neurones within the molecular layers were also labelled as were a significant proportion of the cells in the granular cell layer. In the midbrain, weak labelling was seen in the oculomotor nerve nuclei. The presence of abundant neuromelanin in the substantia nigra pigmented neurones confounded the assessment for dystrophin expression. Strong labelling were noted within the neurones (pontine nuclei) and the surrounding neuropil in the pontine base. In the medulla, the neuropil within the inferior olivary nuclear ribbon showed a strong DYSA signal. Unexpectedly, there was strong labelling of the axons within the vagus and hypoglossal nerves. The respective nuclei displayed dystrophin expression in the surrounding neuropil. The anterior horn cells in the spinal cord showed robust dystrophin expression within the neuronal cell bodies and dendrites (**Fig. 8e**). Of note, these motor neurones expressed both *5’* and *3’* labeled *DMD tr*anscripts (**Fig. 8e-f**).

### Dystrophin and *DMD* expression within the non-neural cell types

In the CS23 choroid plexus, only RNAscope *DMD 3’* labeling was seen, indicative of only short isoforms being expressed (**Fig. 9a**). In contrast, both probes labelled the choroid plexus at 14pcw, but antibody reactivity was clearly detected only with DYS2 (**Fig. 9b**). At CS23, only the RNAscope *DMD 3*’ signal was noted in the blood vessel wall with corresponding exclusive DYS2 immunoreactivity. At 14 pcw stage, the vessel walls showed expression of both RNAscope *DMD 5’* and *DMD 3’* labeled transcripts, and immunoreactivity for both DYSA and DYS2 antibodies. (**Fig. 9e**). In the adult brain, the leptomeningeal, penetrating arteries and arterioles showed predominant expression of RNAscope *DMD 5’* labeled transcripts localised to the vascular smooth muscle cells (**Fig. 9f** and **9g**). At the isoform level, *dp427m* was consistently expressed in the smooth muscle coats of the leptomeningeal and cortical penetrating arteries and arterioles (**Fig. 9f** and **9g**). *Dp71* transcripts were noted in the parenchymal capillary walls and the glia limitans (**Fig. 9h** and **9i**).

**Fig. 9.**
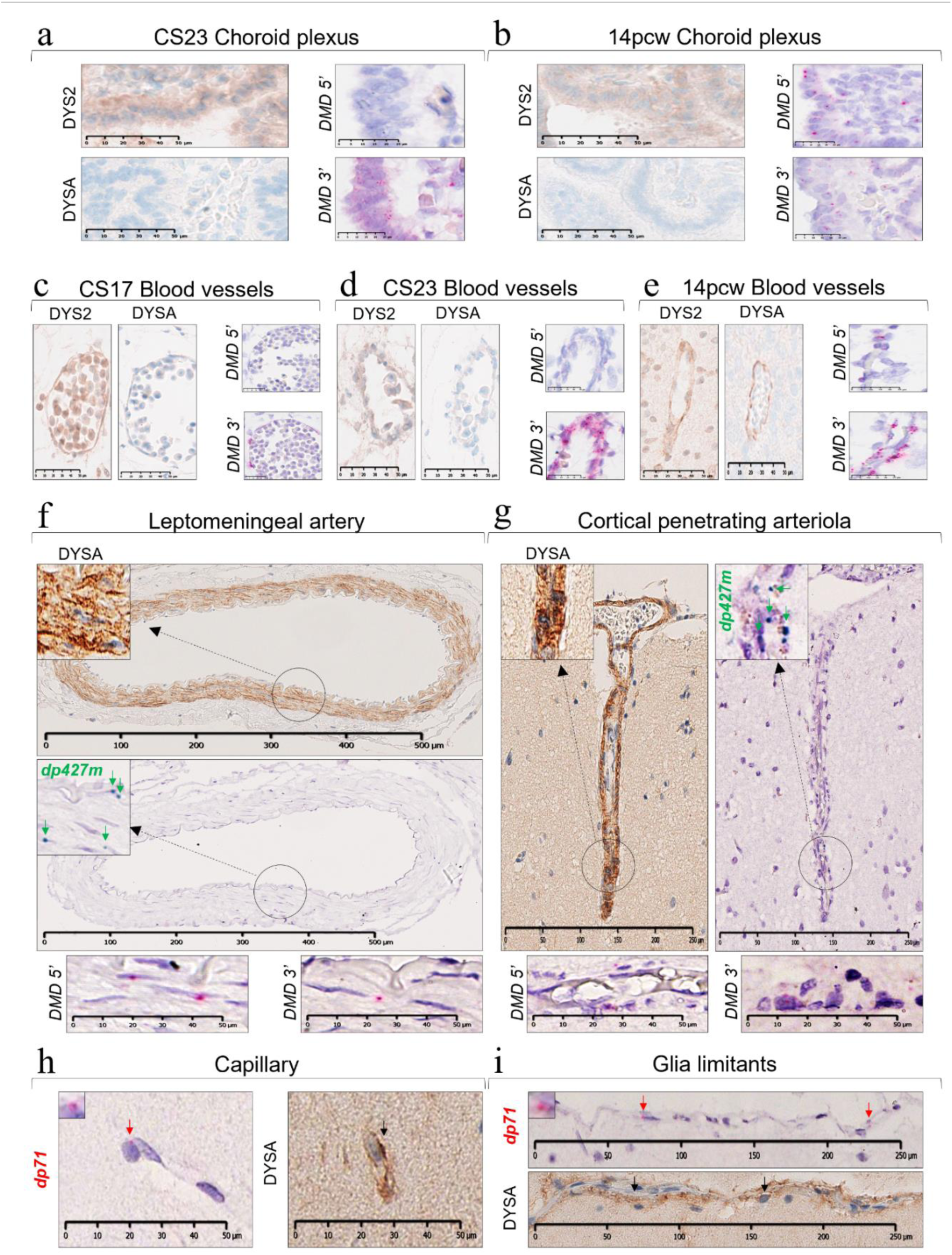
Dystrophin protein and transcript expression in brain vasculature. Embryonic and foetal choroid plexus and blood vessels stained with DYSA and DYS2 antibodies and hybridised with the RNAscope *DMD 5’* and *DMD 3’* probes (**a**-**e**). Adult blood vessels stained with DYSA antibody and processed for RNAscope or Duplex Basescope (**f**-**i**). Note positive signal detected with duplex Basescope (*Dp427m*-green/*Dp71*-red), RNAscope *DMD 5’ DMD 3’* probes and the DYSA antibody within the tunica media of a medium-sized leptomeningeal artery (**f**) and in a cortical penetrating arteriole (**g**) in the ventromedial prefrontal/orbitofrontal cortex. Note also expression of the *Dp71* transcript and DYSA immunoreactivity in the cerebellar cortical parenchymal capillary (**h**) and cerebellar pial-glial membrane (glia limitans) at the cortical surface (**i**). Each red arrow indicates RNAscope or duplex Basescope (*dp71*) signal (red dots) while green arrows indicate duplex Basescope (*dp427m*) signal (green dots).

### Developmental regulation of dystrophin protein isoforms in the human brain assessed by capillary western blot

As immunohistochemical cellular localisation of different dystrophin isoforms in FFPE sections was hindered by the lack of isoform-specific dystrophin antibodies, analysis of protein isoform expression and semi-quantification was carried out by WES using a C-terminal antibody (Ab154168) in protein extracts from frozen developing and adult brains (**Fig. 10**). Protein peaks with the molecular weights of dp427, dp140, and dp71 were detected in all the brain areas from developing and adult samples, while dp116 was detected in 3 out of 5 adult brain areas (cerebellum, amygdala, and dorsolateral prefrontal cortex).

**Fig. 10.**
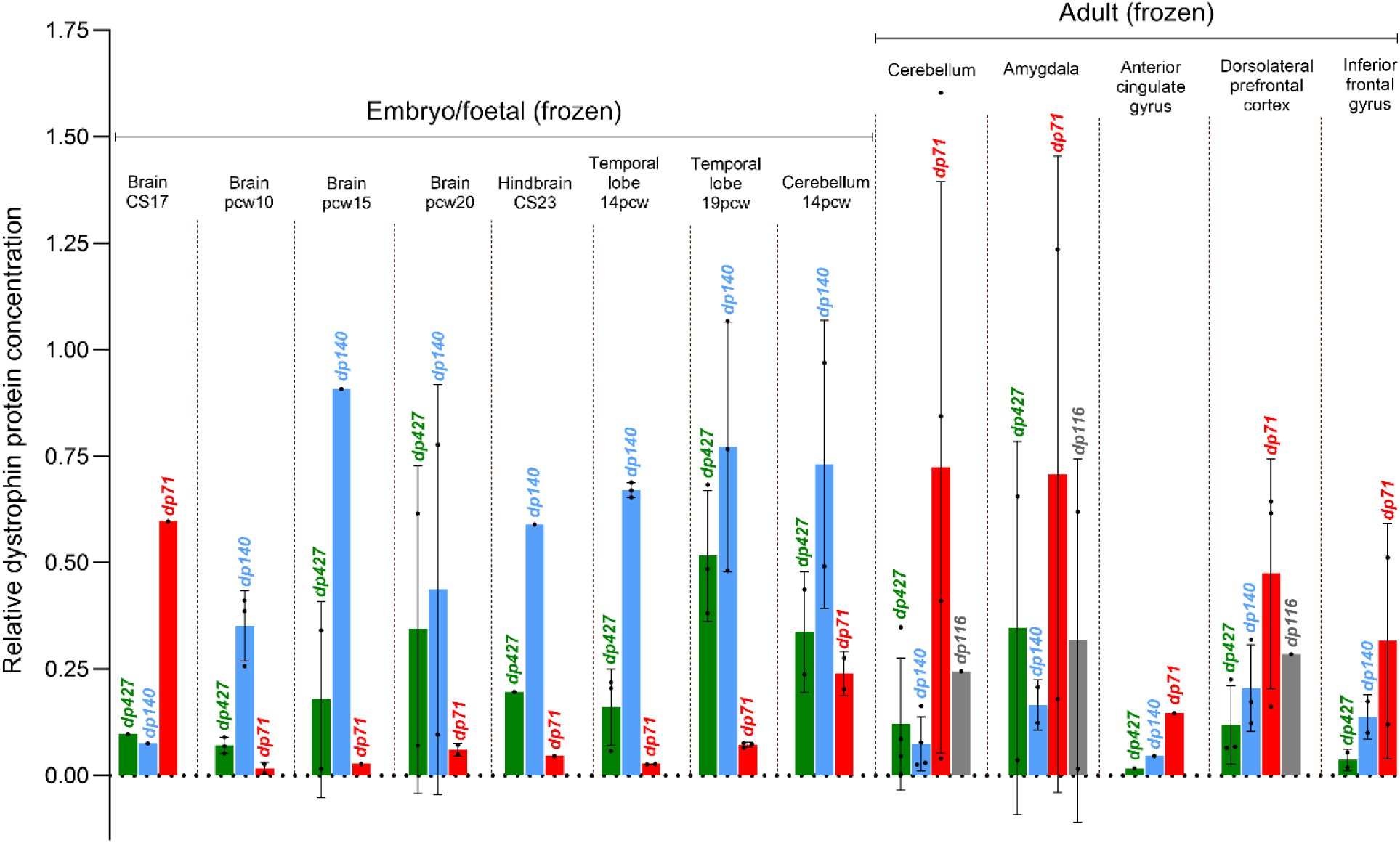
Detection of dystrophin in developing and adult human brain regions by capillary western blot. Relative dystrophin protein concentration (Dp427) in brain/brain regions from embryos (n=2), foetuses (n=15) and adults (n=12) detected by the C-terminal antibody Ab154168. Pcw: postconceptional weeks; CS: Carnegie stage.

During development, apart from CS17, dp140 was the predominant isoform detected in all brain regions assayed with an average relative protein concentration (RPC) of 0.58, followed by dp71 (average RPC=0.25), and dp427 appeared to be the less expressed isoform (average RPC=0.11), with the exception of CS17 brain (RPC=0.6);. In contrast, dp71 was the predominant isoform in all adult brain regions, and dp140, though expressed throughout was greatly reduced as compared to developing brains, as particularly evident in the cerebellum (RPC=0.73 in developing cerebellum versus 0.07 in adult). Dp116 was selectively expressed in adult cerebellum (RPC=0.24), amygdala (average RPC=0.32) and dorsolateral prefrontal cortex (RPC=0.28), but not in cingulate and frontal gyrus nor in embryonic/foetal samples. Although this technique did not allow us to pinpoint which of the long isoforms was expressed, it showed that dp427 was expressed, though to different extents, in all adult brain regions examined.

Together, good concordance between protein and transcript isoform expression was observed, with changes in dp140 and dp71 protein expression from developing to adult brain closely mirroring transcript changes, with the former being higher prenatally and the latter postnatally. However a degree of discordance between the levels of some protein and transcript was noted especially in some adult brain areas, potentially due to non-specific tissue contamination in the frozen specimen pocessed for WES.

## Discussion

This study provides a comprehensive analysis of isoform-specific dystrophin expression in developing and adult pathology-free human brains combining a variety of quantitative methods (RT-qPCR and immunoblot), *in situ* hybridisation and immunohistochemistry techniques that deliver, for the first time, a rich neuroanatomical, cellular and sub-cellular context to the dystrophin expression profile.

Our RNAscope studies, that allow for spatial localisation of the *DMD* transcripts, provide novel information on the complexities of *DMD* expression in the human. The *DMD 5’* signal appeared as large juxtanuclear/intranuclear dots, suggestive of transcriptional ‘hotspots’ containing multiple full-length nascent and mature transcripts, whereas the *DMD 3’* signal consisted of multiple small cytoplasmic dots, indicating the presence of mature transcripts as previously described in the dog by Hildyard et al., [39].

Our results show that in the human brain *DMD 3’* signal was robust throughout the embryonic and foetal developmental stages, whereas the *DMD 5’* signal increased from early to late development. Remarkably, *DMD* transcripts were already abundant in neuroepithelial cells at an early developmental stage, CS17 (6 weeks of gestation). The dystrophin interacting protein, dystroglycan, has been suggested to play a role in the maturation of the mouse perinatal neural stem cell niche, partly via regulation of Notch signalling, an important pathway in neural development [40, 41]; therefore, it is tempting to speculate that dystrophin may modulate Notch signalling in the human neuroepithelium.

Furthermore, the hippocampus and cerebellar Purkinje cell layer also showed high expression of *DMD* transcripts. The complete absence of *DMD 5’* signal in the granule cells of the developing and adult cerebellum was striking, suggestive of a cell-specific specialised role for the shorter isoforms. Altogether, our RNAscope data showed that *DMD* transcripts are ubiquitously expressed in the developing and adult human brain.

When isoform-specific *DMD* transcripts were spatially mapped by Basescope and quantified by RT-qPCR, *dp427c* showed consistent high expression across most regions in the developing and the adult brain. This is in contrast to earlier reports showing low-level expression across human brain development [10]. Also in contrast to previous findings reporting no expression of *dp427p* in the developing human brain [10, 42], we found this isoform highly expressed (RT-qPCR) in the embryonic and foetal brains. Like *dp427p,* expression of *Dp427m* was high during development, but more variable in the adult brain. Most of the transcripts were expressed in the leptomeningeal and/or the parenchymal blood vessels within the smooth muscles. Unexpectedly, as not previously reported, consistent expression of *dp427m*, though at very low-levels, was observed in neurons and oligodendroglial-like cells across different brain regions.

*Dp140* was highly expressed across all brain regions in the embryonic and foetal brain. In the adult brain its expression was significantly downregulated, with sparse to few transcripts noted in the cerebellum (Purkinje cells, granule cells and dentate neurons), hippocampus (dentate granule cells and hilar neurons), amygdala and across all neocortical areas (mostly in pyramidal neurons). Similarly, prior investigations reported a developmental regulation of *dp140* characterised by high transcripts expression in foetal samples which remain stable in cerebral cortex and cerebellum in adults [10]. Our findings also support the hypothesis that the *dp140* isoform is pivotal during development and it persists in the adult brain in areas as previously proposed [10, 39] and likely to undergo adult neurogenesis, such as the subgranular zone in the dentate gyrus of the hippocampus.

In keeping with earlier findings, *dp71/dp40* expression was consistently high during development and persisted in the adult brains across all regions. Transcripts were identified within neurons, parenchymal capillary walls (likely astroglia), and the glia limitans. Low-to-moderate *dp40* expression was noted throughout development and persisted in the adult brain.

In addition, we have demonstrated for the first time co-expression of two different *DMD* isoforms within the same neuron by duplex Basescope assays. This can be observed from the early developmental stages to adulthood across all anatomical brain regions. However, neurons co-expressing *DMD* isoforms represent less than ca. 2% of the total neuronal population.

It should be noted that gene co-expression studies suggest distinct roles for *dp427*, *dp140* and *dp71*-related dystrophin signalling in the brain. Our study has also shown that the granule cells of the dentate gyrus in the hippocampus, neurons in the pontine nuclei, and the cerebellar granular cells displayed the most diversity in *DMD* isoform expression.

With regard to protein expression our western blot results showing good correlation with the RT-qPCR data, confirmed for the first time that the dp140 protein was the predominant isoform expressed in embryonic brains, whereas dp71 was more abundant in adult brains.

Dp427 was consistently expressed throughout development and in adult life. Unexpectedly, we detected expression of dp116 both at the transcripts and protein level in developing and adult brains, mainly in the cerebellum and dorsolateral prefrontal cortex. This indicates that dp116 expression may not be restricted to the Schwann cells as previously suggested [10]. We also showed by RT-qPCR that *dp260-1* is the only *DMD* isoform not expressed in the CNS.

Furthermore, to support the substantial presence of dystrophin in the brain, immunolabeling with the DYSA antibody showed dystrophin expression mainly in the somata and proximal dendrites of a diverse population of neurons, including pyramidal cells in the neocortex and hippocampus, granule cells of the dentate gyrus, Purkinje cells, granule cells and dentate neurons in the cerebellum, and neuronal populations in the deep grey nuclei (basal ganglia and thalamus), brain stem and cervical cord (anterior horn cells).

The use of duplex RNAscope assays allowed us to study cellular coexpression of dystrophin and markers of GABAergic (*GAD1)* and glutamatergic (*SLC17A7)* neurons circumventing limitations of other approaches (e.g. poor reactivity of dystrophin antibodies in FFPE brain sections and lack of isoform-specific dystrophin antibodies), and to demonstrate dystrophin expression in both *GAD1+* inhibitory interneurons and *SLC17A7+* excitatory neurons in all neocortical regions, cerebellum and the hippocampus. Furthermore, the extensive co-expression of *GAD1+* and *DMD 3’* transcripts observed in the 14pcw developing Purkinje layer suggest that intermediate and short-length dystrophins might play a crucial role in the cerebellar GABAergic circuitry maturation. Our data also demonstrate a switch from these isoforms to full length dystrophin in adult GAD1+ Purkinje neurons, as indicated by significant expression of *DMD 5’* transcripts in these neurones. Transient expression of non-full length isoforms in GAD1+ Purkinje neurons is interesting, as *Dp71*-null mice display alterated glutamatergic transmission and cognitive deficits [43], and is consistent with a realatively early onset of developmental defects leading to at least some of the neural deficit observed in DMD patients.

The dystrophin isoform brain localisation reported here provides an important underpinning to better understand the common brain involvement in DMD patients.

Notably, although impacted by limitations mainly related to small sample size, clinical studies showed that proximal and distal DMD mutations are linked to type and severity of different brain co-morbidities [11, 44, 45].

In most patients, distal mutations are associated with more severe intellectual disabilities mainly linked to the loss of *dp140* and *dp71* [11, 46–48] leading to reduced intelligent quotients and working memory index scores [11] along with poorer academic performances [49].

Contrarily, proximal mutations affecting *dp427* have been associated with neuropsychiatric manifestations ranging from an increased startle response, indicative of enhanced anxiety and authistic spectrum and attention deficit hyperactivity disorders [11, 12, 50].

It has been speculated that the lack of dp427 might be compensated by shorter dystrophins including dp140 and dp71, hence their absence exacerbates the cognitive and neuropsychiatric phenotypes [11]. In this context, our data showing a strong *dp427, dp140* and *dp71* expression in developing and adult brains support this hyphothesis.

Our findings on dystrophin expression in brain blood vessels both during development and in adults are consitent with its previously reported expression in muscle blood vessels; interestingly, impaired angiogenesis was observed in muscle of DMD patients and animal models [51–54]. In addition, consistent with a possible vascular phenotype in DMD patient brain is the reduced perfusion observed in the *mdx* mouse brain [55] and impaired cerebral blood flow in DMD patients demonstrated by magnetic resonance imaging studies [56].

We are aware that our study has a number of limitations. Although we presented a temporally resolved dystrophin expression profile, the numbers in each age-group are very small. Information regarding dystrophin expression in the crucial post-natal and paediatric age-group is missing, because pathology-free brains for assessment in this age group were not available. Inherent variability (biological and technical) within and in between samples must be taken in to account while interpreting the data. In particular, accurate quantification of the *in situ* hybridisation signal was hampered by the sheer number and complexity of neuronal and glial elements in tissue sections, further compounded by the lack of clear cell boundaries, and often saturation of the particulate chromogenic signal within individual neurons.

Given the rapid advancement of single-cell and single-nucleus RNA sequencing technologies as well as single cell proteomics, it will eventually be possible to unravel the complexity of the brain dystrophin cellulome/proteome/interactome. However, validation of such findings in human pathology-free brains as well as DMD brains will remain challenging due to the aforementioned reasons.

## Conclusions

In summary, we have shown that *DMD* and dystrophin isoforms are abundantly expressed in the human brain across the lifespan and are developmentally regulated, and localised them at single-cell resolution. We have demonstrated for the first time co-expression of different *DMD* isoforms across several key brain areas in the developing and adult human brain at the regional and single-cell level.

We also have shown for the first time ubiquitous expression of dystrophin within different glutamatergic neuronal populations in the developing and adult human brain. This finding should prompt future studies investigating the molecular pathology of dystrophin signalling at the neuronal excitatory synapse. In addition, we have provided novel information on *DMD* gene expression in GABAergic and glutamatergic neuronal populations in the human brain. The granular localisation of dystrophin reported here will also underpin therapeutic developments aimed at restoring dystrophin functionality in the CNS, hence reducing the burden of brain comorbidities and improving the quality of life of DMD patients.

## Acknowledgements

The support of Sarepta Therapeutics for providing financial support for developing this research project. The authors would also like to thank the NIHR GOSH biomedical research centre (BRC) for providing part of the facilities and equipment and both the Human Developmental Biology Resource (http://hdbr.org) and the Edinburgh Brain Bank as part of BRAIN UK network for providing the human specimens. Special thanks are extended also to the General Department of Health Affairs at Princess Nourah bint Abdulrahman University and Saudi Arabian Cultural Bureau and Ministry of Education (scholarship to RA).

Finally, a special acknowledgement goes to Dr Karl Frontzek and Dr Maria Thom for providing annotations of a set of brain images and to Jenifer Morgan and Federica Montanaro for their support and useful discussions throughout the project.

## SUPPLEMENTARY INFORMATION

**Supplementary Fig. 1.**
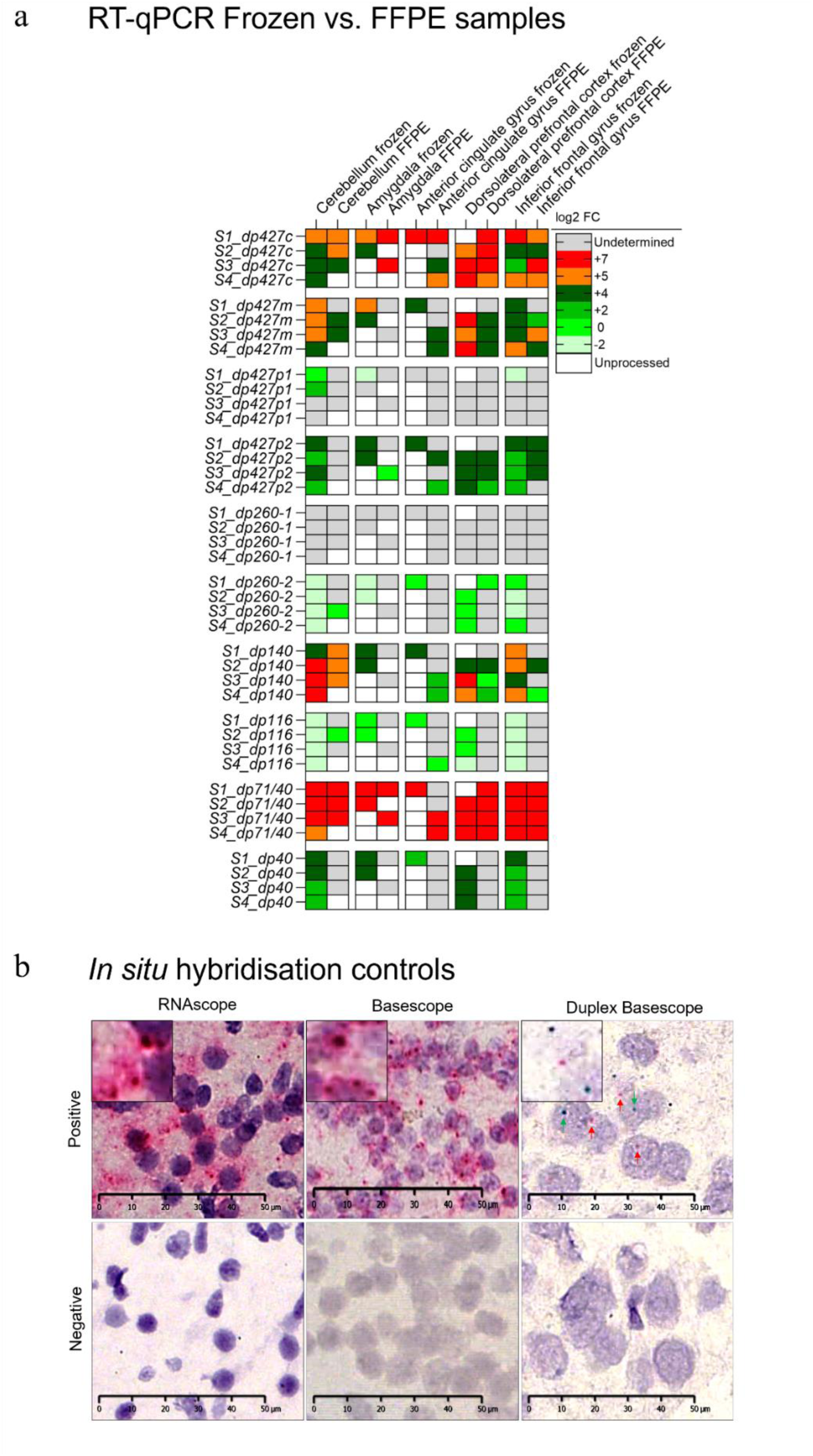
Expression patterns of the main *DMD* transcripts in matching FFPE/frozen adult brain areas assessed by RT-qPCR (**a**). Expression is normalised to the brain specific *RPL13* housekeeping gene. Colour code: white: unprocessed; light green: low expression; green: medium expression; orange/red: high expression; grey: undetermined. FFPEs: Formalin Fixed Paraffin Embedded tissue samples. Representative images showing the *in situ* hybridisation results of control sections incubated with the RNAscope/Basescope *UBC* (ubiquitin C) and PBS and/or probe diluent (negative) (**b**).

**Supplementary Fig. 2.**
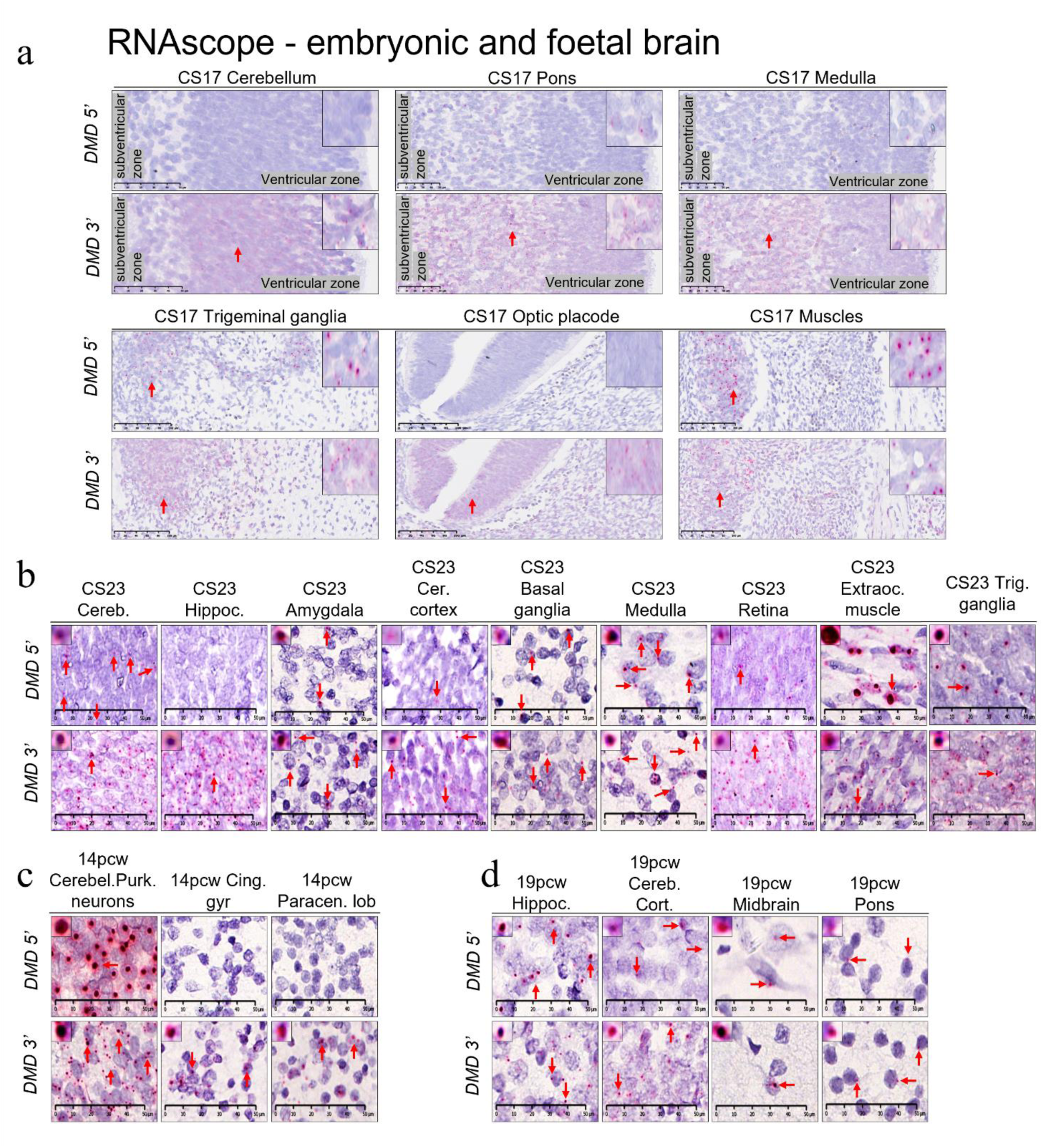
Representative images of CS17 embryonic (**a**), embryonic CS23 (**b**), foetal 14pcw (**c**) and foetal 19 pcw (**d**) areas hybridised with either RNAscope *DMD 5’* or *DMD 3’* probes (counterstained with haematoxylin) showing stable red signal. Each red arrow indicates RNAscope signal (red dots) or representative dots in case of high RNA expression/clusters. The top left panel shows the most representative puncta to a higher magnification. Pcw: postconceptional weeks; CS: Carnegie stage. Scale bars = 25 and 50 µm.

**Supplementary Fig. 3.**
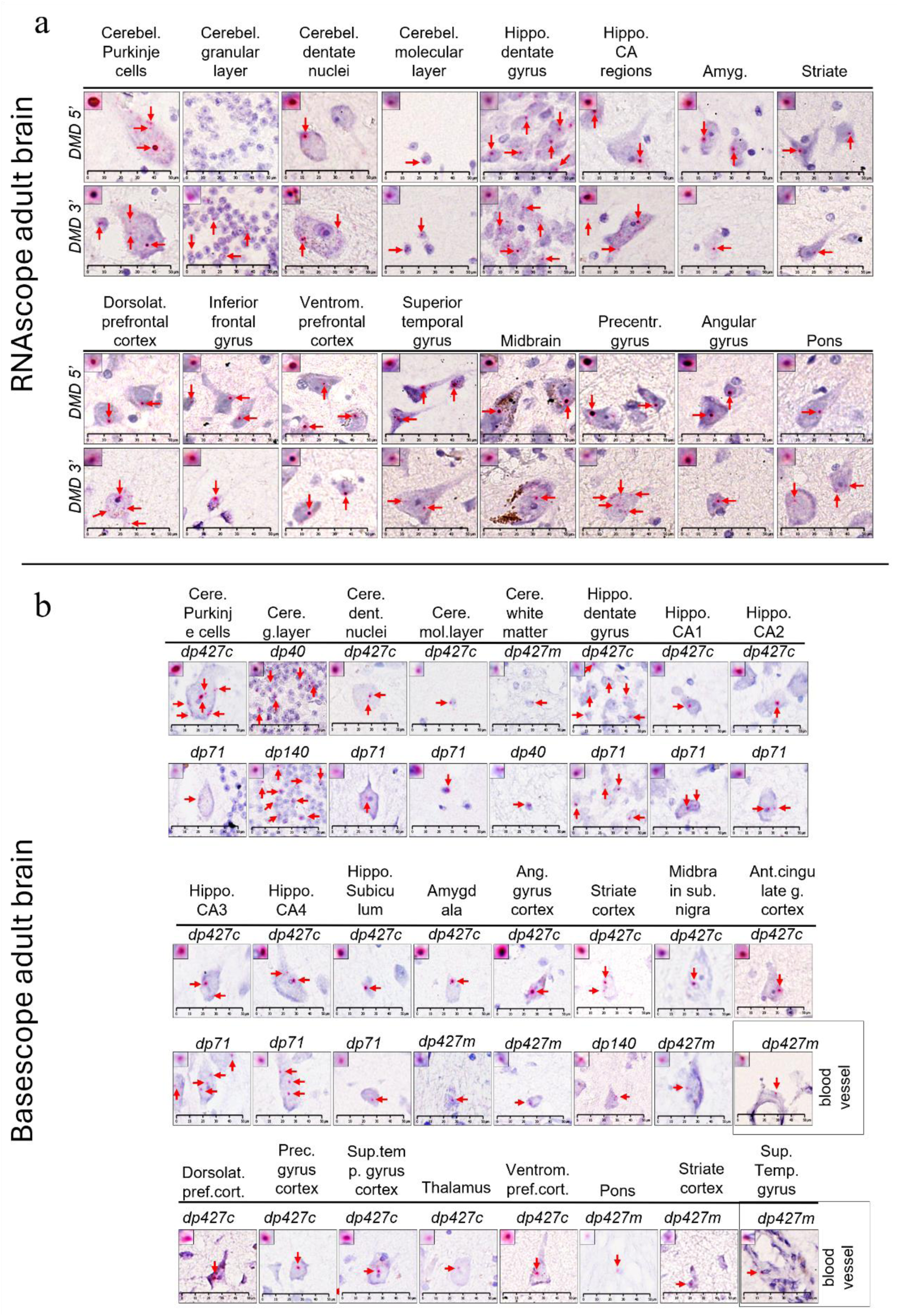
*In situ* hybridisation experiments in adult human brain areas. Representative images of adult brain areas hybridised with either RNAscope *DMD 5’* or *DMD 3’* probes (**a**) or singleplex Basescope probes (**b**). The top left panel shows the most representative puncta to a higher magnification. Each red arrow indicates RNAscope/Basescope signal (red dots) or representative dots in case of high RNA expression/clusters. Hippocampus CA: Cornu Ammonis. Scale bars = 50 µm.

**Supplementary Fig. 4.**
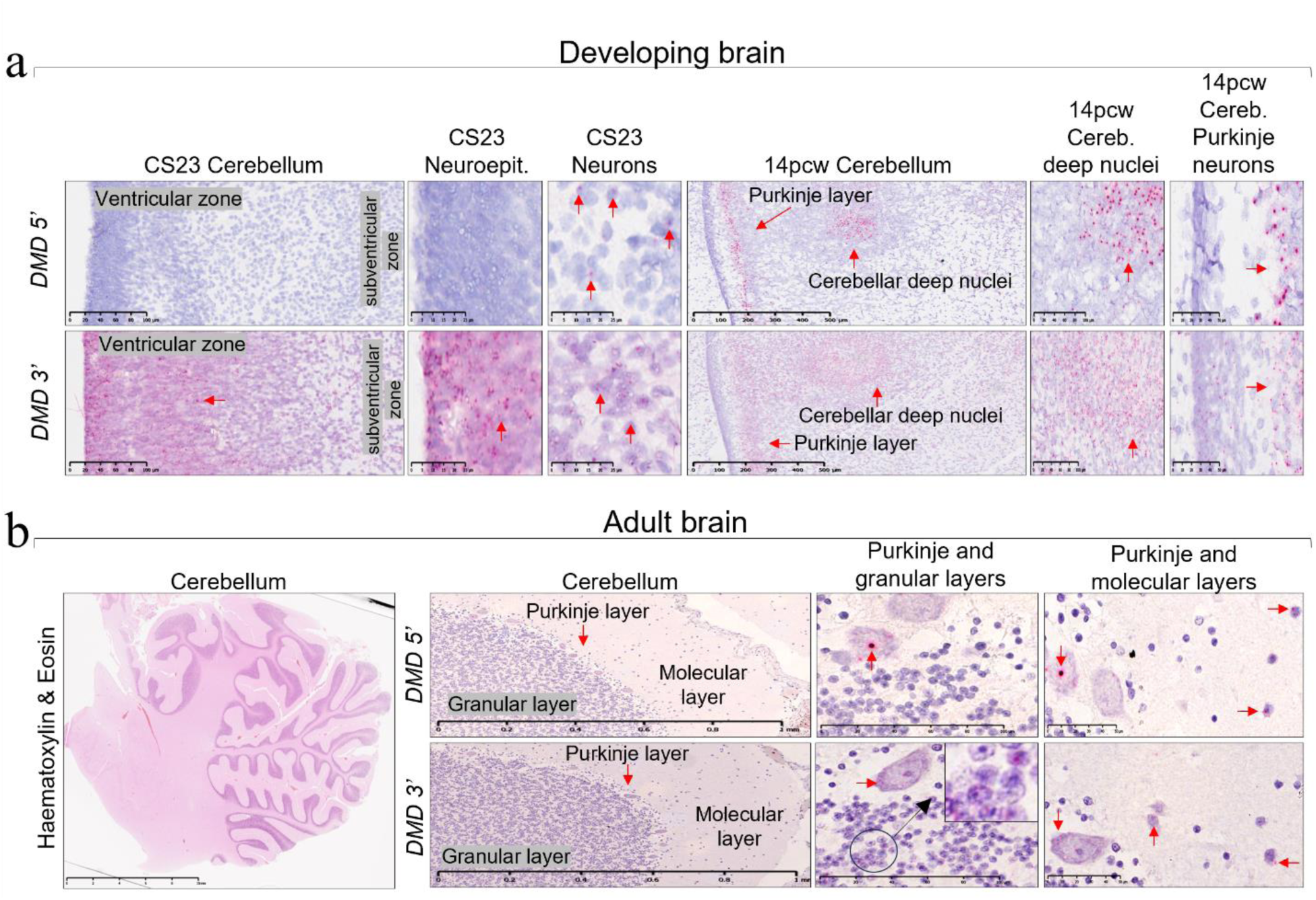
Developmental regulation of *DMD 5’* and *DMD 3’* ends in the human cerebellum. Representative images showing the RNAscope signal in embryonic, foetal (**a**) and adult cerebellum (**b**). Each red arrow indicates RNAscope signal (red dots) or representative dots in case of high RNA expression/clusters. Pcw: postconceptional weeks; CS: Carnegie stage.

**Supplementary Fig. 5.**
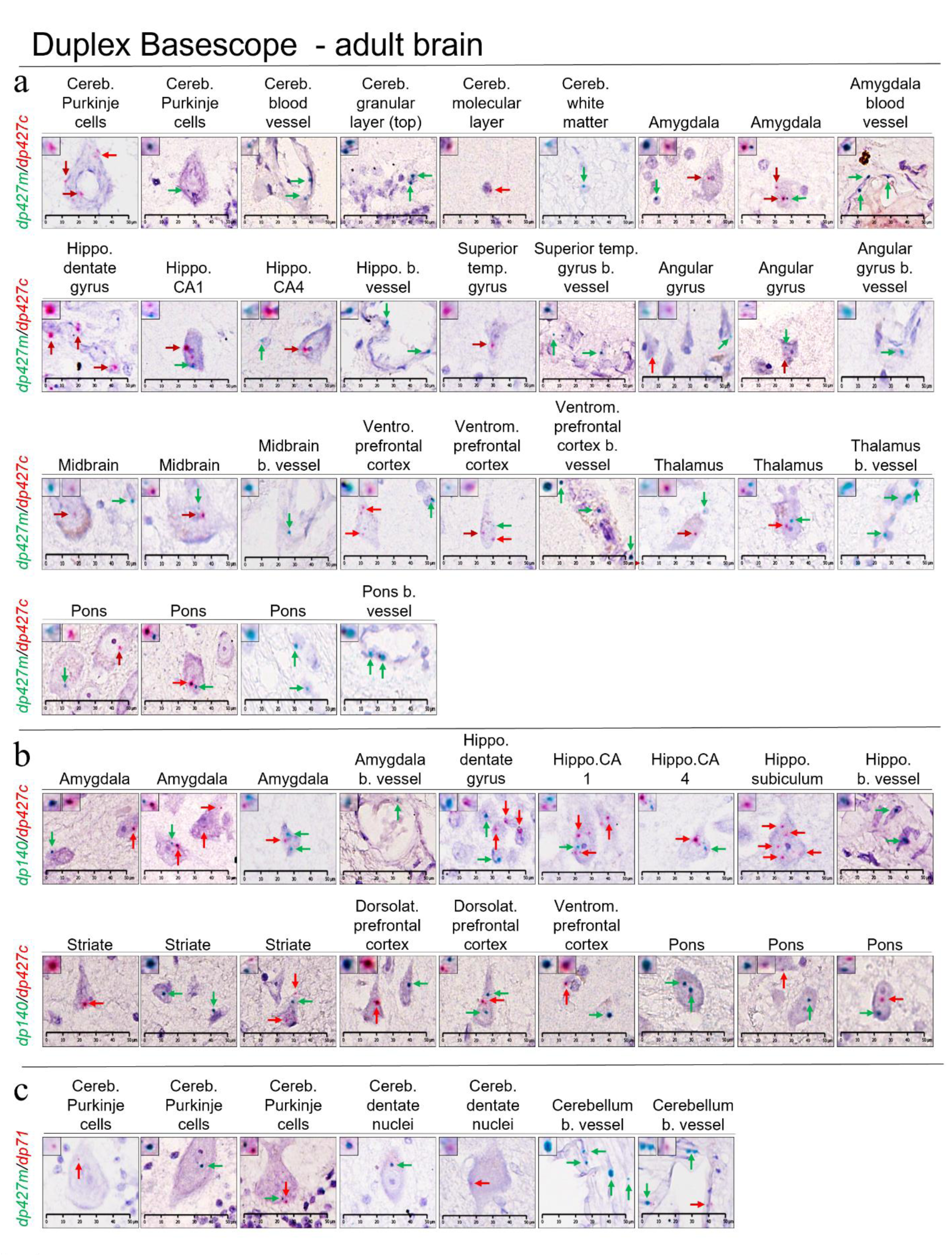
Duplex Basescope results from adult FFPE brain sections. Representative image of the results from the adult samples processed for four different combinations: *dp427m*(green)/*dp427c*(red) (**a**), *dp140*(green)/*dp427c*(red) (**b**) and *dp427m*(green)/*dp71*(red) (**c**). Red and green arrows indicate RNA transcripts (dots). The top left inset shows the largest puncta to a higher magnification. Hippocampus CA: Cornu Ammonis. Scale bars = 50 µm.

**Supplementary Fig. 6.**
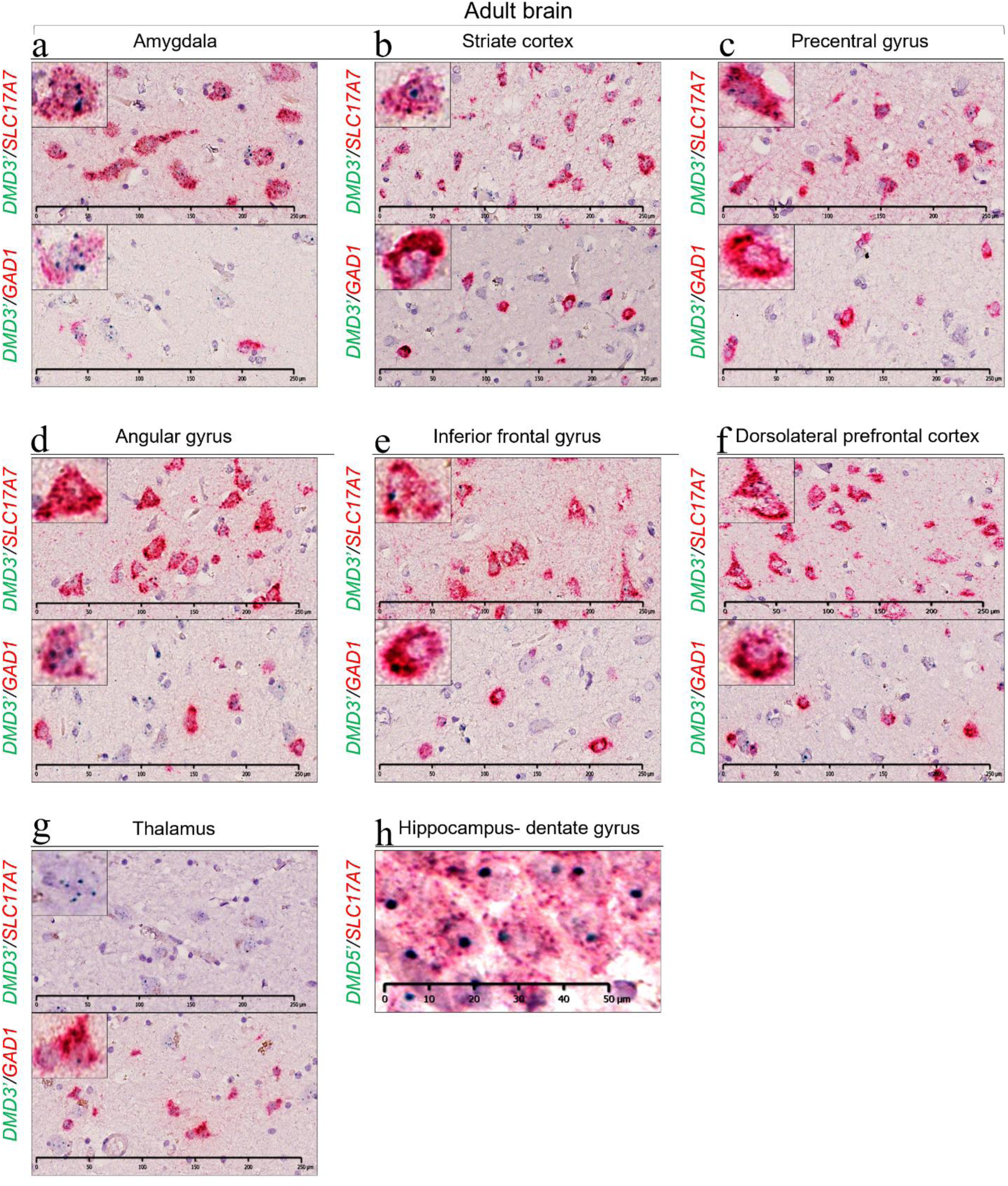
Duplex RNAscope results from adult FFPE brain sections. Representative images of the results from the adult samples (**a**-**h**). Green dots indicate RNAscope *DMD 5’* and *DMD 3’* transcripts. Red puncta indicate *GAD1* (GABAergic) or *SLC17A7* (glutamatergic) transcripts.

**Supplementary Table 1.** Details on donors and corresponding specimens used in the study. DOD: date of death. Pcw: postconceptional weeks; CS: Carnegie stage. NA: not available; d: days; w: weeks; S1-4: subject.

| **Adults - Frozen and Formalin-fixed paraffin-embedded (FFPEs)** |  |  |  |  |  |  |
| --- | --- | --- | --- | --- | --- | --- |
| **Subject** | **Brain Bank N - ID for publication** | **Gender** | **Age** | **DOD-processed\*** | **Primary cause of death** | **Histopathological features (brain)** |
| S1 | SD022/16 - BBN001.28959 | Male | 39 y | 7 days | Ischaemic heart disease | No significant abnormalities |
| S2 | SD025/13 - BBN_15223 | Male | 47 y | 8 days | Coronary artery atherosclerosis | No significant abnormalities |
| S3 | SD034/14 - BBN_22628 | Male | 49 y | 8 days | Presumed cardiac arrhythmia | No significant abnormalities |
| S4 | SD053/14 - BBN_24342 | Male | 33 y | 5 days | Ischaemic heart disease | No significant abnormalities |
| *FFPEs only |  |  |  |  |  |  |
| **Adults - Matched FFPE and frozen specimens from the same donor** |  |  |  |  |  |  |
| S1 | SD022/16 - BBN001.28959 | Cerebellum, amygdala, anterior cingulate gyrus, inferior frontal gyrus |  |  |  |  |
| S2 | SD025/13 - BBN_15223 | Cerebellum, amygdala, dorsolateral prefrontal cortex, inferior frontal gyrus |  |  |  |  |
| S3 | SD034/14 - BBN_22628 | Cerebellum, dorsolateral prefrontal cortex, inferior frontal gyrus |  |  |  |  |
| S4 | SD053/14 - BBN_24342 | Cerebellum, dorsolateral prefrontal cortex, inferior frontal gyrus |  |  |  |  |

| **Embryonic/foetal - Frozen** |  |  |  |  |  |  |
| --- | --- | --- | --- | --- | --- | --- |
| **Subject** | **Brain Bank N - HDBR ID** | **Gender** | **Age** | **DOD-processed** | **Regions analysed** | **Histopathological features (brain)** |
| S1 | CS17 - 10840 | NA | 41 d | NA | Brain | NA |
| S2 | CS17 - 13715 | NA | 41 d | NA | Brain | NA |
| S1 | 10 pcw - 10967 | NA | 12 w | NA | Brain | NA |
| S2 | 10 pcw - 10897 | NA | 12 w | NA | Brain | NA |
| S3 | 10 pcw - 11816 | NA | 12 w | NA | Brain | NA |
| S1 | 15 pcw - 14212 | NA | 17 w | NA | Brain | NA |
| S2 | 15 pcw - 14105 | NA | 17 w | NA | Brain | NA |
| S3 | 15 pcw - 13587 | NA | 17 w | NA | Brain | NA |
| S1 | 20 pcw - 13886 | NA | 22 w | NA | Brain | NA |
| S2 | 20 pcw - 13589 | NA | 22 w | NA | Brain | NA |
| S3 | 20 pcw - 13988 | NA | 22 w | NA | Brain | NA |
| S1 | CS23 - 14760 | NA | 56 d | NA | Cerebellum | NA |
| S2 | CS23 - 15002 | NA | 56 d | NA | Cerebellum | NA |
| S1 | 14 pcw - 16323 | NA | 16 w | NA | Cerebellum | NA |
| S2 | 14 pcw - 16092 | NA | 16 w | NA | Cerebellum | NA |
| S3 | 14 pcw - 15999 | NA | 16 w | NA | Cerebellum | NA |
| S1 | 14 pcw - 15658 | NA | 16 w | NA | Temporal lobe | NA |
| S2 | 14 pcw - 13927 | NA | 16 w | NA | Temporal lobe | NA |
| S3 | 14 pcw - 15095 | NA | 16 w | NA | Temporal lobe | NA |
| S1 | 19 pcw - 13782 | NA | 21 w | NA | Temporal lobe | NA |
| S2 | 19 pcw - 13885 | NA | 21 w | NA | Temporal lobe | NA |
| S3 | 19 pcw - 14160 | NA | 21 w | NA | Temporal lobe | NA |
| S1 | CS17 - 15484 | NA | 41d | NA | Hindbrain | NA |
| S1 | CS23 - 12388 | NA | 56 d | NA | Hindbrain | NA |
| S2 | CS23 - 12391 | NA | 56 d | NA | Hindbrain | NA |
| S3 | CS23 - 12193 | NA | 56 d | NA | Hindbrain | NA |
| S1 | CS23 - 14452 | NA | 56 d | NA | Medulla | NA |

**Supplementary Table 1.** Details on donors and corresponding specimens used in the study.
| **Embryonic/foetal - FFPEs** |  |  |  |  |  |  |
| --- | --- | --- | --- | --- | --- | --- |
| **Subject** | **Brain Bank N - HDBR ID** | **Gender** | **Age** | **DOD-processed** | **Regions analysed** | **Histopathological features (brain)** |
| S1 | CS17 - 12242 | NA |  | NA | Head | NA |
| S2 | CS23 - 13141 | NA |  | NA | Head | NA |
| S1 | 14pcw-15295 | NA |  | NA | Hindbrain | NA |
| S2 | 14pcw-15658 | NA |  | NA | Hindbrain | NA |
| S3 | 14pcw-13883 | NA |  | NA | Temporal lobe | NA |
| S1 | 14pcw-15095 | NA |  | NA | Temporal lobe | NA |
| S2 | 19pcw-15699 | NA |  | NA | Temporal lobe | NA |
| S3 | 19pcw-14211 | NA |  | NA | Temporal lobe | NA |
| S1 | 19pcw-15026 | NA |  | NA | Temporal lobe | NA |
| S2 | 19pcw-12519 | NA |  | NA | Cerebellum, Pons | NA |

**Supplementary Tab. 2.** Specifics of the probes used for the ISH experiments.

| Ref | Probe | Comments |
| --- | --- | --- |
| 720291 | BaseScope Probe - BA-Hs-DMD-Dp427c | recognises *dp427c* |
| 720301 | BaseScope Probe - BA-Hs-DMD-Dp427m | recognises *dp427m* |
| 720321 | BaseScope Probe- BA-Hs-DMD-Dp427p2 | recognises *dp427p2* |
| 720281 | BaseScope Probe - BA-Hs-DMD-Dp140 | recognises *dp140* |
| 720331 | BaseScope Probe - BA-Hs-DMD-Dp71 | recognises *dp71* |
| 1105461 | BaseScope Probe- BA-Hs-DMD-Dp40 | recognises *dp40* |
| 712121 | BaseScope Probe - BA-Hs-UBC-1zz | recognises *UBC* (Ubiquitin C) |
| 71071 | BaseScope Probe - BA-Hs-PPIB-1zz | recognises *PPIB* (Peptidylprolyl Isomerase B) |
| 459561 | RNAscope 5' Probe - Hs-DMD-Dp427m-E2-E10 - NM_004006.2 | recognises *dp427* only (c, m and p) |
| 1039481 | RNAscope 3' Probe - Hs-DMD-Dp427m-E63-E70 - NM_004006.2 | recognises all dystrophin isoforms |
| 310041 | RNAscope Positive Control Probe- Hs-UBC | recognises *UBC* (Ubiquitin C) |
| 404031-C2 | RNAscope Probe - Hs-GAD1 | recognises *GAD1* (glutamate decarboxylase 1) |
| 415611-C2 | RNAscope Probe - Hs-SLC17A7 | recognises *SLC17A7* (solute carrier family 17 vesicular glutamate) |
| Ref | Kits |  |
| 322360 | RNAscope 2.5 HD Detection Reagents |  |
| 322500 | RNAscope 2.5 HD Duplex Detection Reagents kit |  |
| 323900 | BaseScope Reagent Kit v2 – RED Assay |  |
| 323800 | BaseScope Duplex Reagent Kit |  |
| 310091 | RNAscope Wash buffer reagents |  |
| 322000 | RNAscope pretreatment reagents |  |

**Supplementary Tab. 3.**
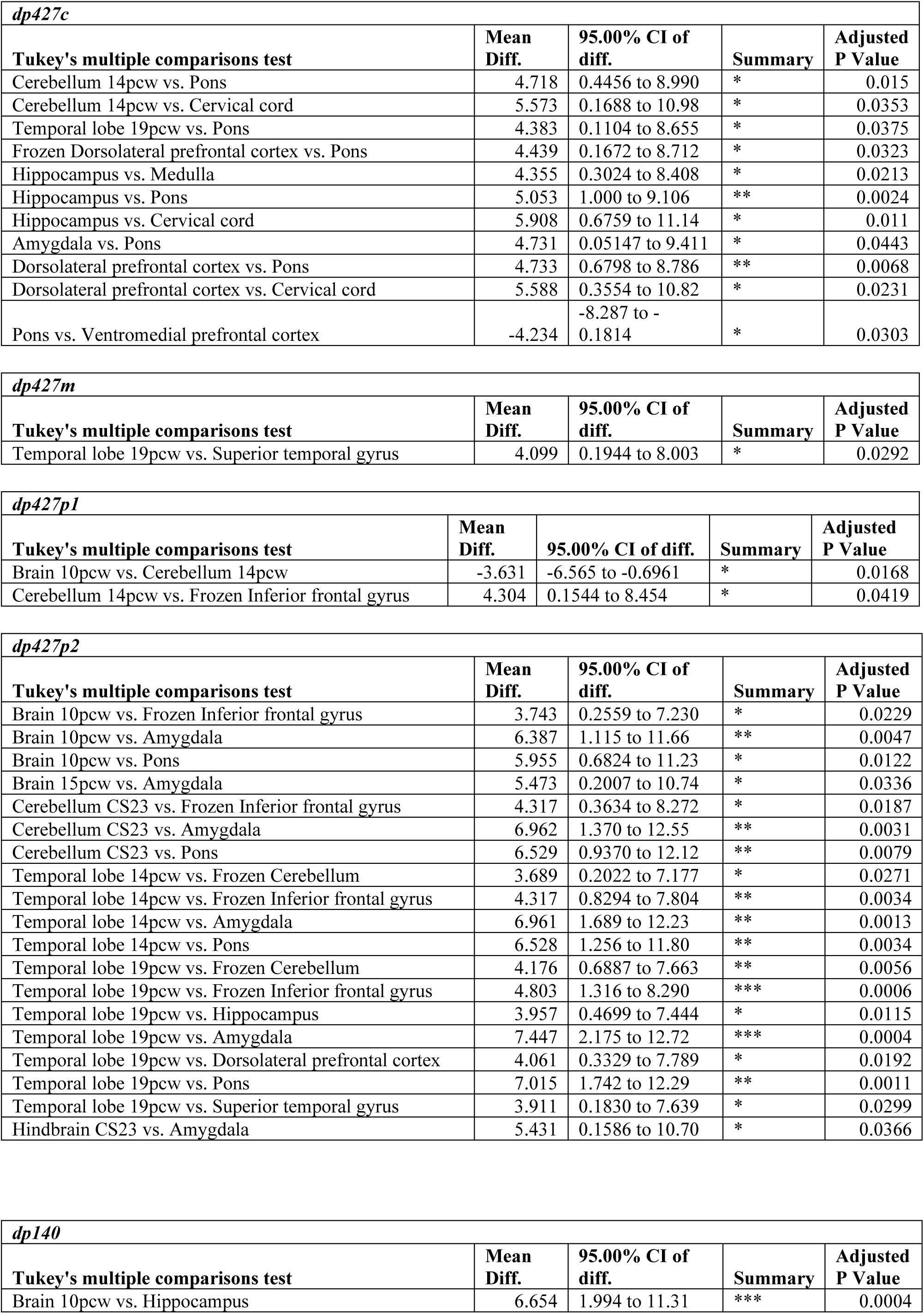
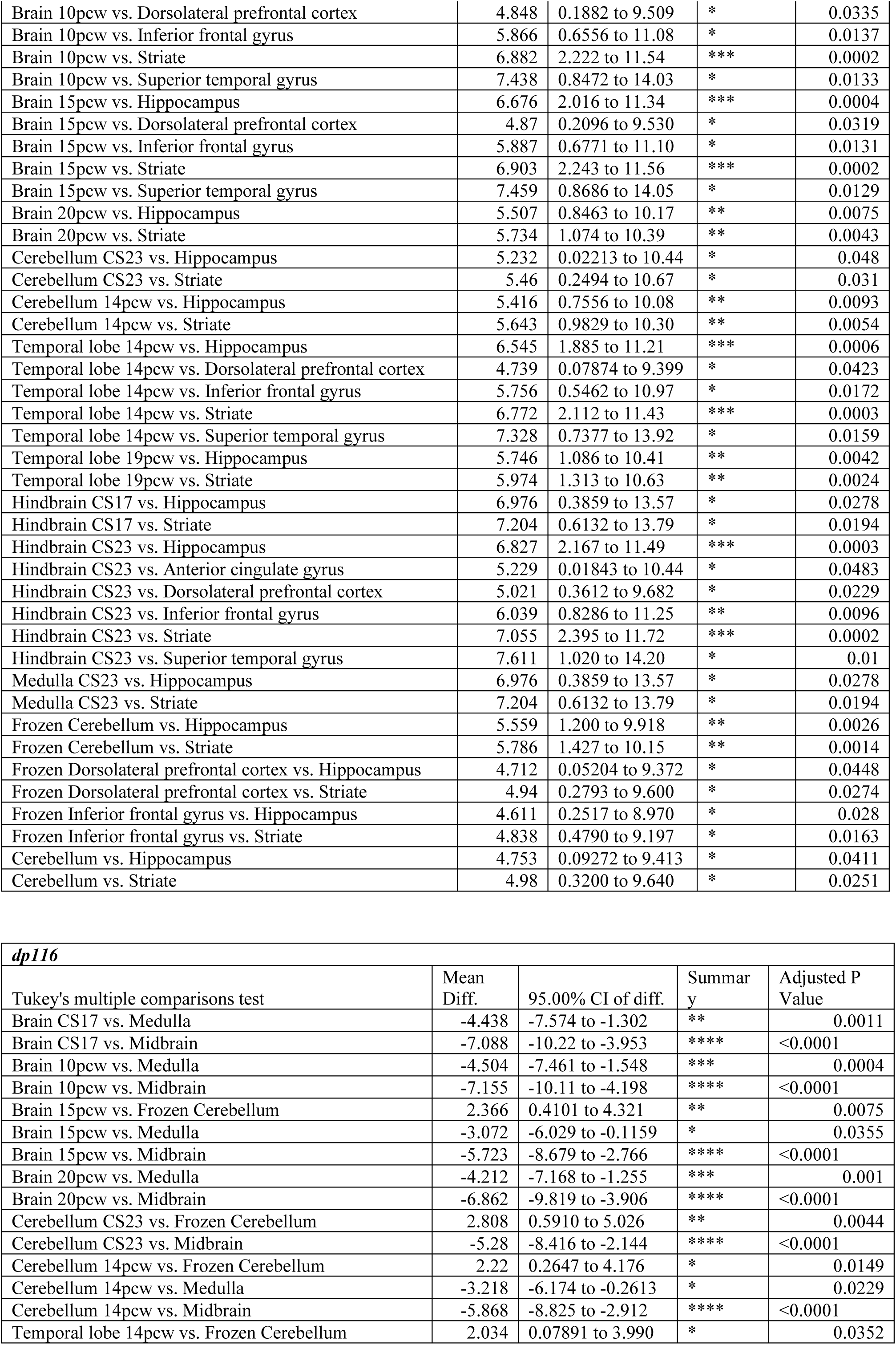
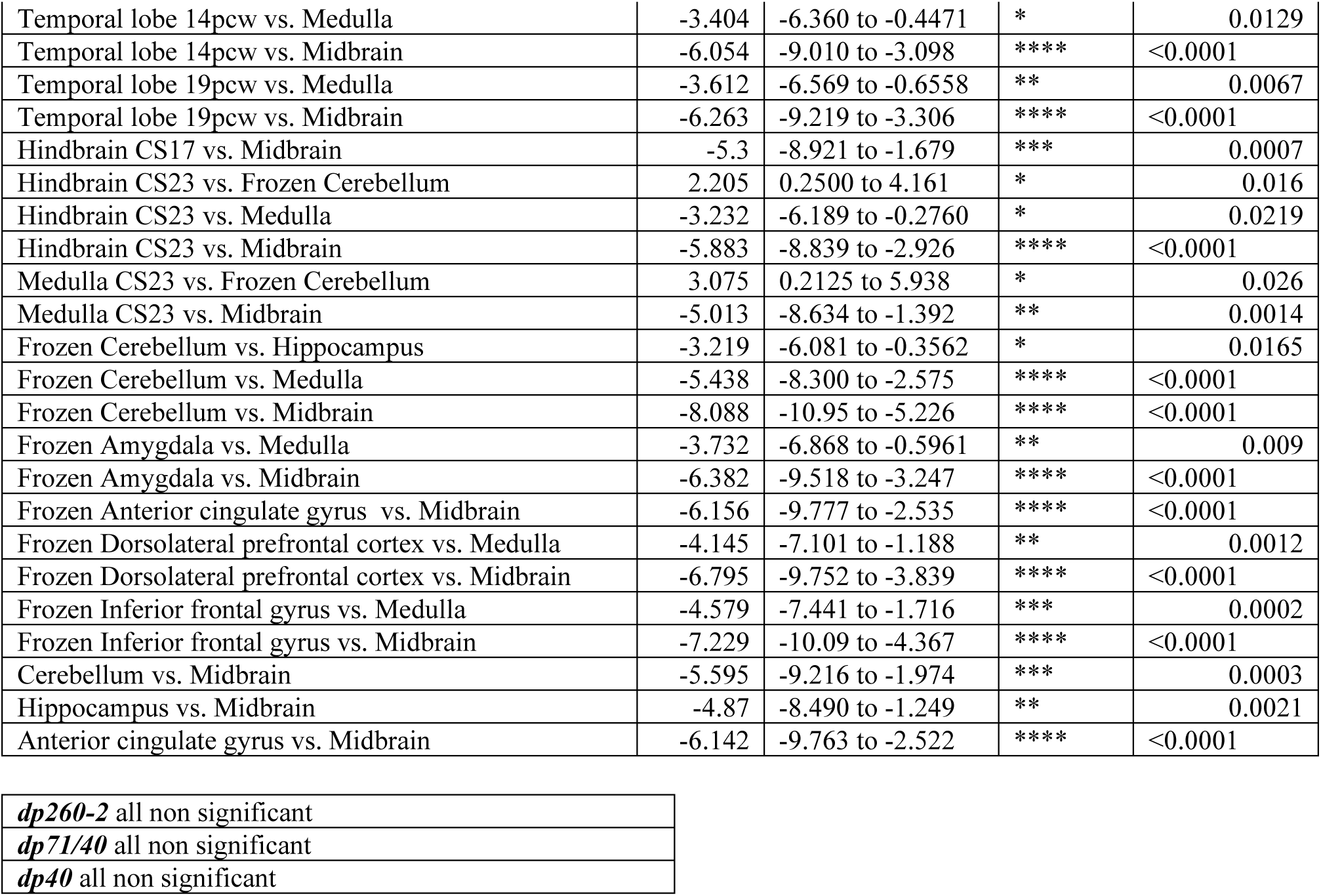
Statistical analysis of isoform specific-*DMD* RT-qPCR data.

